# Lessons from a multilaboratorial task force for diagnosis of a fatal toxoplasmosis outbreak in captive primates in Brazil

**DOI:** 10.1101/2023.09.27.559677

**Authors:** Francine Bittencourt Schiffler, Silvia Bahadian Moreira, Asheley Henrique Barbosa Pereira, Igor Falco Arruda, Filipe Romero Rebello Moreira, Mirela D’arc, Ingra Morales Claro, Thalita de Abreu Pissinatti, Liliane Tavares de Faria Cavalcante, Thamiris dos Santos Miranda, Matheus Augusto Calvano Cosentino, Renata Carvalho de Oliveira, Jorlan Fernandes, Matheus Ribeiro da Silva Assis, Jonathan Gonçalves de Oliveira, Thayssa Alves Coelho da Silva, Rafael Mello Galliez, Debora Souza Faffe, Jaqueline Goes de Jesus, Marise Sobreira Bezerra da Silva, Matheus Filgueira Bezerra, Orlando da Costa Ferreira Junior, Amilcar Tanuri, Terezinha Marta Castiñeiras, Renato Santana Aguiar, Nuno Rodrigues Faria, Alzira Paiva de Almeida, Alcides Pissinatti, Ester Cerdeira Sabino, Maria Regina Reis Amendoeira, Elba Regina Sampaio de Lemos, Daniel Guimarães Ubiali, André F.A. Santos

## Abstract

As exemplified by the Coronavirus Disease 2019 (COVID-19) pandemic, infectious diseases may emerge and spread rapidly, often causing serious economic losses and public health concerns. In fact, disease outbreaks have become increasingly common, especially those of zoonotic origin. The Brazilian Ministry of Health is responsible for national epizootic surveillance. However, the system’s focus primarily on diseases affecting humans has led to the neglect of other zoonotic diseases. In this report, we present an integrated investigation of an outbreak that occurred during the first year of the COVID-19 pandemic among captive neotropical primates housed at a primatology center in Brazil. After presenting a range of non-specific clinical signs, including fever, prostration, inappetence, and abdominal pain, ten primates from five different species died within approximately four days. Despite the state of health emergency due to the pandemic, a network of volunteer researchers was established to investigate the outbreak. A wide range of high-resolution techniques was used for different pathogens, including SARS-CoV-2 (RTq-PCR, ELISA and IHC), Toxoplasma gondii (IHC and IFA) and Escherichia coli (IFA), as well as a portable Metagenomic Sequencing utilizing Nanopore Technology. Within a span of four days after necropsies, we successfully identified T. gondii as the causative agent of this outbreak. This case highlights some of the obstacles faced with the current Brazilian surveillance system, which is still limited. A cross-platform interdisciplinary investigation could be a possible model for future epizootic investigations in non-human animals.

**Author summary:** The Brazilian epizootic surveillance system, under the regulation of the Ministry of Health, has been established to address a national list of compulsory notifiable diseases. However, focusing mainly on the risks to humans causes other zoonoses to be neglected. Here we present an outbreak that occurred during the first year of the COVID-19 pandemic that affected eleven neotropical primates (NP) belonging to six different species. Within four days of exhibiting a range of non-specific clinical signs, including fever, prostration, inappetence, and abdominal pain, ten NPs died. Despite testing negative for pathogens included in the national surveillance policy, a collaborative group of researchers investigated the outbreak in detail. Using integrated diagnostic techniques, we identified *Toxoplasma gondii* as the causative agent four days after necropsy. Toxoplasmosis causes devastating acute death outbreaks in neotropical primates and is currently absent in the national guidelines. This unified effort proved the effectiveness of a multidisciplinary collaborative surveillance network in facilitating precise diagnoses.

## 1. Introduction

Between September and November 2020, during the first year of the Coronavirus Disease 2019 (COVID-19) pandemic and concomitant with the emergence of the P.2 strain in Brazil [1], ten neotropical primates (NP) from five different species died within approximately four days of the onset of clinical signs at the *Centro de Primatologia do Rio de Janeiro* (CPRJ) in Guapimirim, Rio de Janeiro, Brazil. Thus, in view of the reduced staff due to the mandatory quarantine and the official count of 141,406 deaths to that date, a health alert was issued to deal with the situation.

In Brazil, epizootic surveillance is overseen by the Brazilian Ministry of Health through the National Surveillance System. To facilitate data integration, the information system called *Sistema de Informação de Agravos de Notificação* (SINAN) was implemented in 2006, along with the establishment of a national list of compulsory notifiable animal diseases. This list includes rabies, plague, influenza, and several arboviruses, such as yellow fever, Oropouche, Mayaro, West Nile and equine encephalomyelitis viruses [2,3]. Thus, when an animal is found dead or sick with any of the diseases on this list, it is mandatory to report the incidence to the nearest health department [4,5].

While the notification is formally required for all classes of animals, the majority of the reports are related to NP, which are well-known sentinels for yellow fever virus (YFV) epizootics. In order to prevent the transmission of YFV to humans, most surveillance guidelines are tailored towards monitoring NPs and mosquitoes [5]. In recent decades, there has also been an increasing concern regarding the rabies virus (RABV), primarily due to the rise in the number of cases observed in non-traditional reservoirs, such as NPs [6]. Consequently, when a NP dies, the standard procedure is to send samples to national surveillance reference laboratories specializing in YFV and RABV. However, if samples test negative for the compulsory notifiable diseases, the investigation is typically discontinued.

The COVID-19 pandemic evidenced how infectious diseases may emerge suddenly and become a serious threat to global health [7]. First cases were confirmed in China in early December 2019 [8] and only two months later Brazil already registered its first case [9], reaching 194,949 deaths by the end of 2020. Several other important zoonotic viruses have also jumped from non-human animals to humans causing human epidemics in the last 15 years. These resulted in the emergence of other coronaviruses such as SARS-CoV, and the Middle East respiratory syndrome–related coronavirus (MERS-CoV) [8,10], influenza viruses, including H1N1 [11], filoviruses such as Ebola and Marburg viruses; [12,13], and arboviruses including Zika, chikungunya, yellow fever viruses [14,15]. Analogous to human epidemics, epizootics are infectious diseases affecting a considerable number of nonhuman animals at the same time and region. For instance, large epizootics caused by foot-and-mouth disease virus and by the ongoing spread of highly pathogenic avian influenza (H5N1; [16,17]) can have severe socioeconomic consequences and often require coordinated public health approaches between countries [18,19].

Here, we present an integrated epidemiological and genomic investigation of this outbreak among captive NPs at the CPRJ/Brazil. A couple days after necropsies, we identified that toxoplasmosis was the cause of the deaths. It is an important disease for neotropical primate medicine due to the fast progression and the large number of hosts affected. It is important to highlight that the CPRJ serves as a scientific breeding facility for endangered NPs and is located in a rural area adjacent to the *Três Picos* State Park, encompassing approximately 160,000 acres of Atlantic Forest and housing 380 specimens from 29 species of NP (**S1 Fig)**. Despite being in a national state of health emergency due to the pandemic, we mobilized a collaborative interdisciplinary network comprising research and public health laboratories to conduct a comprehensive investigation of the outbreak, including NP and staff, testing for various pathogens using a wide range of high-resolution diagnostic techniques. In addition, professionals exposed to these animals underwent clinical and laboratorial examination and received instructions on the procedures to be followed in case of an acute febrile illness.

## 2. Materials and methods

### 2.1. Ethics statement and Official Notification

All procedures conducted in this study adhered to the biosafety requirements established by the World Health Organization (WHO). Ethical approval for the study was obtained from the Ethics Committee on the Use of Animals in Scientific Experimentation at the Health Sciences Center from the *Universidade Federal do Rio de Janeiro* (CEUA-CCS/UFRJ) under protocol number: 066/20. The collection and transportation of samples were further approved by the Biodiversity Authorization and Information System (SISBIO) of the *Instituto Chico Mendes de Conservação da Biodiversidade* (ICMBIO) (license number 75941-1). In addition, the present study received approval by the local ethics review committee at the *Hospital Universitário Clementino Fraga Filho* (*Certificado de Apresentação de Apreciação Ética* – CAAE: 30161620.0.0000.5257) and by the national ethical review board (CAAE: 30127020.0.0000.0068) for the collection of biological samples and testing of CPRJ workers. Regarding human sample collection, all participants included in the study were adults aged 18 years or older and provided consent through an informed consent form. CPRJ workers were tested for SARS-CoV-2 and *Yersinia sp.*, as described below, and for Hepatitis A, B, C (HAV, HBV, HCV), Cytomegalovirus (CMV), toxoplasmosis and leptospirosis through the Brazilian Unified Health System. All information obtained during the outbreak investigation were made available to the Rio de Janeiro State Health Secretary through detailed reports elaborate throughout the outbreak.

### 2.2. Animals

Our study focused on monitoring an infectious disease outbreak that took place at the CPRJ between September and November 2020, resulting in the death of 11 captive NPs. During this period, all deceased primates underwent necropsy, including two individuals of *Brachyteles arachnoides*, three *Alouatta ululata*, two *A. guariba clamitans*, one *A. caraya*, two *Cacajao melanocephalus*, and one *Plecturocebus caligatus*. All procedures conducted during the necropsies were carried out in full compliance with and approved by the Brazilian Ministry of the Environment (SISBIO 30939–12).

### 2.3. Sample Collection for Molecular Testing and Genetic Material Extraction

Since clinical signs appeared almost simultaneously in different NPs species, the investigation was carried out in parallel in nine different laboratories, each employing different diagnostic methods until the pathogen was definitely identified. It should be noted that the order in which these methods were applied may vary.

A variety of samples were analyzed in this study, including blood and swab samples obtained from multiple specimens at different time points during the outbreak, as well as tissue fragments collected during necropsies. Blood samples from the femoral and saphenous veins of NP (preferably) and by peripheral venipuncture from CPRJ staff, were collected in EDTA tubes. For serological analysis, serum was isolated by centrifugation at 400 g for 10 min. Swab samples were obtained by inserting sterile swabs into the selected cavity (rectal or oral for NPs, and nasopharyngeal for humans), rotated slightly, allowing for a 10-second period to absorb secretions, and then removed with slow circular movements. Swabs were stored with 1 mL of RNAlater^®^ (Invitrogen, Thermo Fisher Scientific, Waltham, MA, USA). Specimens were temporarily stored at room temperature and permanently stored at –80 °C.

Total nucleic acid extraction from swab and tissue samples was performed using ReliaPrep™ Viral TNA Miniprep System (Promega, Madison, WI, USA). Swab samples were simply inverted in the tube and briefly centrifuged prior to extraction. Tissue extraction involved disrupting approximately 1 mm^3^ with 500 μL RNAlater^®^ solution using Lysing Matrix E (MP Biomedicals do Brasil, São Caetano do Sul, SP, BR) on a Super FastPrep-2 (MP Biomedicals, Valiant, CN) through 30-second cycles alternating with an ice bath until complete dissolution. The mixture was then centrifuged at 6,500 g at 4 °C, and 200 μL of the supernatant was used, following manufacturer’s protocol. Nucleic acid extraction from blood samples was performed using the MasterPure^™^ Complete DNA and RNA Purification Kit (Lucigen, LGC Ltd, Teddington, GB). For metagenomic analysis, samples were centrifuged for 5 min at 10,000 g before extraction.

### 2.4. Necropsy and histopathology

As the animals died, they were subjected to necropsy by the veterinary pathologists at the *Setor de Anatomia Patológica* from *Universidade Federal Rural do Rio de Janeiro* (SAP/UFRuralRJ), together with members of the CPRJ and the *Serviço de Criação de Primatas Não-humanos* from *Instituto de Ciência e Tecnologia em Biomodelos* (SCPrim ICTB). A total of 11 NP underwent the procedure, which was performed using personal protective equipment (PPE) compatible with Biosafety Level-3 (BSL-3), including specific lab coats, gloves and eye protection. During necropsies, organs (skin, brain, lymph nodes, lung, heart, trachea, esophagus, thyroid, adrenal, spleen, kidney, liver, stomach, and intestines) were collected in 10% buffered formalin and fixed 24-48h for routine histological processing. Fragments of approximately 1 cm^3^ were also excised and stored with 1 mL of RNAlater^®^ until nucleic acid extraction. Standardized brain 5-section trimming was performed, corresponding to the structures: telencephalon, hippocampus, amygdaloid nuclei, diencephalon, mesencephalon, ventricular system, cornu ammonis, cerebellum, pons, and myelencephalon. Sections were stained with hematoxylin and eosin (HE) for optical microscopy.

### 2.5. Detection of Viral Agents

#### 2.5.1. SARS-CoV-2

All procedures were performed by the *Laboratório de Diversidade e Doenças Virais* (LDDV), the *Laboratório de Virologia Molecular* (LVM) and the *Núcleo de Enfrentamento e Estudos de Doenças Infecciosas Emergentes e Reemergentes* (NEEDIER) from UFRJ. Genetic material extracted from nasopharyngeal swabs from CPRJ workers and oral swabs and tissues (lung, trachea and intestine) from NP were used for molecular detection in a One Step Reverse Transcription-quantitative Polymerase Chain Reaction (RT-qPCR) system, using GoTaq^®^ Probe qPCR Master Mix (Promega, Madison, WI, USA) and the CDC 2019-Novel Coronavirus (2019-nCoV) Real-Time RT-PCR Diagnostic Panel (Integrated DNA Technologies, Coralville, IA, USA), according to the manufacturer’s instructions. This protocol targets the SARS-CoV-2 N1 and N2 genes and, as an internal control, the human ribonuclease P (RNaseP) gene. All reactions were performed in a 7500 Real-Time PCR System (Applied Biosystems, Waltham, MA, USA). Samples were considered positive when both targets (N1 and N2) amplified with cycle threshold (Ct) ≤ 37.

For serological diagnosis, an Enzyme-Linked Immunosorbent Assay (ELISA) protocol was performed for both staff members from CPRJ and NP. The 96-well ELISA plates (Corning, USA) were coated overnight at 4 °C with 200 ng per well of the SARS-CoV-2 protein S produced by the *Laboratório de Engenharia de Cultivos Celulares* (LECC) from the UFRJ by Prof. Leda Castilho [20]. After a cycle of five washes with phosphate-buffered saline (PBS) 0.05% Tween-20, the plates were blocked with 100 µL of 5% Bovine Serum Albumin (BSA) and incubated for 2 h at room temperature. Each serum sample was tested at a dilution of 1:50 in PBS 0.05% Tween-20 with 2% BSA and Bromocresol Purple, added to the wells and incubated for 1 h at 37 °C. After a new cycle of five washes, 50 μL of horseradish peroxidase (HRP)-conjugated goat anti-Monkey IgG (1:10,000, Thermo Fisher Scientific, USA) was added to each well, the plate was incubated at 37 °C for 1 h, followed by a new cycle of washes. Lastly, 50 μL of TMB substrate (3,3’,5,5;-tetramethylbenzidine; Thermo Fisher Scientific, USA) was added into each well. After 10 min incubation, the reaction was stopped by adding 50 μL of 1 M H_2_SO_4_ solution. Reactions were analyzed at 450 nm wavelength and results were expressed in optical density (OD). All samples were tested in duplicate and, therefore, were considered reactive when the mean of the ODs of the replicates exceeded the cut-off, obtained by calculating the mean of the ODs of the negative controls plus three times the standard deviation. To normalize the OD values between reaction plates, the percentage of the difference between the OD of the well with sample and the average OD of the white wells (only with the protein) was calculated.

To evaluate the presence of coronaviruses other than SARS-CoV-2, samples of the lungs and trachea of necropsied NPs, as well as blood samples from sick NPs were used for PCR amplification with a set of PanCoronavirus (PanCov) primer sequences (α-, β-, γ-, and δ-coronaviruses)[21]. After RNA extraction, synthesis of cDNA was performed with the High-Capacity cDNA Reverse Transcription Kit (Thermo Fisher Scientific, Waltham, MA, USA), followed by PCR reaction with Platinum™ Taq DNA Polymerase (Invitrogen, Thermo Fisher Scientific, Waltham, MA, USA). This reaction consisted in 2.5 μL of 10X PCR Buffer (1X), 1 μL of 50 mM MgCl_2_ (2 mM), 0.2 μL of 25 mM dNTP (0.2 mM), 1 μL of each primer (150 pmol), 0.25 μL Platinum™ Taq DNA Polymerase (1.25 U), 5 μL cDNA template and 15 μL Nuclease-free Water. The reaction was conducted with an initial activation at 94 °C for 2 min, followed by 35 cycles of amplification (30 sec at 94 °C, 5 min at 52 °C, 1 min at 72 °C) and a final extension step at 72 °C for 1 min. Results were visualized with a 1% agarose gel electrophoresis.

The Panbio COVID-19 Ag Rapid Test Device (Abbott Rapid Diagnostic Jena GmbH, Jena, TH, DE) was also used to detect the viral nucleocapsid protein in nasopharyngeal samples from CPRJ workers. Detection was performed immediately after sampling, following the manufacturer’s instructions (reading up to 15 min). For NP testing, only oral samples were used, since the small size of the animals prevented the collection of nasal and nasopharyngeal samples. Lastly, to completely rule out a SARS-CoV-2 infection in NP, veterinarians from the SAP/UFRuralRJ further tested lung samples of necropsied NPs by Immunohistochemistry (IHC) using an Anti-Sars-CoV Nucleocapsid Protein (Novus Biologicals^®^, Centennial, CO, catalog no. NB100-56576) [22].

#### 2.5.2. Arenavirus and Hantavirus

Serum samples of NPs were screened for IgG antibodies against recombinant nucleoprotein protein (rN) of *Orthohantavirus andesense* (*Mammantavirinae: Hantaviridae*) and *Mammarenavirus choriomeningitidis* (LCMV; *Arenaviridae*) whole proteins by an ELISA protocol, as previously described [23,24] by the *Laboratório de Hantaviroses e Rickettsioses (LHR)* from *Instituto Oswaldo Cruz* (IOC) *-Fiocruz*.

To evaluate the presence of arenavirus and hantavirus RNA, RNA from blood and tissue samples from NPs were used for RT-PCR amplification using SuperScript™ IV One-Step RT-PCR System kit (Invitrogen, Thermo Fisher Scientific, Waltham, MA, USA) with a set of primers sequences targeting hantavirus nucleoprotein and polymerase genes [25,26], arenavirus glycoprotein [27] and lymphocytic choriomeningitis virus (LCMV) polymerase gene [28]. Results were visualized in a 1.5% agarose gel electrophoresis.

#### 2.5.3. Arboviruses

Blood samples from NPs were also screened for arboviruses in the State Reference Laboratory. After RNA extraction, samples were subjected to a RT-PCR as previously described [29], which targets the highly conserved 5′-noncoding region (5′-NC) of the yellow-fever virus (YFV) genome. Samples were considered positive when presenting a threshold cycle value ≤ 37. Furthermore, the presence of other main circulating arboviruses was evaluated by RT-qPCR using GoTaq^®^ 1-Step RT-qPCR System (Promega, Madison, WI, USA). Specific diagnosis primers were used for Zika [30], dengue [31], chikungunya [32], Mayaro [33], West Nile [34] and Oropouche [35] viruses. Reactions were assembled following the manufacturer’s instructions and were performed in a 7500 Real-Time PCR System (Applied Biosystems, USA).

### 2.6. Metagenomic sequencing

To investigate the presence of pathogens in an unbiased fashion, we performed rapid metagenomic sequencing using portable Oxford Nanopore sequencing Technology. A total of 20 NPs samples from four different tissues were selected for metagenomic sequencing primarily based on histopathological alterations (liver and intestine, n=9), followed by organs from the respiratory tract (due to the suspicion of respiratory virus infection) (lung, n=4). For those for which no tissue sample was collected, we sequenced only blood samples (n=7). Each sample was analyzed separately in a different sequencing library.

After extraction, isolated RNA was transported to the *Instituto de Medicina Tropical* at *Universidade de São Paulo* (IMT/USP), where sequencing libraries were prepared and sequenced following the SMART-9N protocol [36]. Raw FAST5 files were basecalled using Guppy software version 2.2.7 GPU basecaller (Oxford Nanopore Technologies, Oxford, Oxon, GB), then demultiplexed using Guppy barcoder. We performed taxonomic classification using Kraken v. 2.0.7_beta [37] with the miniKraken_v2 database. Interactive visualization plots were generated with Krona v. 2.8.1 [38]. Afterwards, barcoded FASTQ files were mapped to reference genomes using MiniMap2 [39]. Tablet v. 1.19.05.28 [40] was used to visualize mapping files, count mapped reads, and calculate the percentage of genome coverage and sequencing depth.

### 2.7. Bacterial Agents

#### 2.7.1. Yersinia pestis

Because the location where the outbreak occurred overlaps with a known bubonic plague foci [41], sera from the subjects were evaluated for the presence of the *Yersinia pestis*-specific F1 capsular antigen pestis antibodies. For confirmatory purposes, two distinct multi-specie approaches were used: hemagglutination and ELISA-Protein A. The serology tests were performed by the Plague National Reference Service at the *Instituto Aggeu Magalhães* (IAM) *-Fiocruz* and the detailed protocol is described elsewhere [42].

To evaluate the presence of *Y. pestis*, bone marrow and sera samples from all NPs were submitted to bacterial culture. Furthermore, the spleen, lungs, kidney and whole blood of one individual (*A. caraya*, ID 2576) were also submitted to bacterial culture. After sample collection, the cleared phalange bones were sent in a sterile tube to the BSL-3 laboratory in the IAM. The bone was sprayed with ethanol 70% at the external parts and the head of the bone was removed. The bone marrow was collected with a syringe, diluted 1:1 parts in sterile saline solution and with a bacteriological loop, and was transferred to the following media: blood agar base (BAB), agar MacConkey and agar Salmonella-Shigella. The plates were incubated at least for 48h at 28°C and the colonies were tested using the *Y. pestis*-specific bacteriophage-lysis test [43], multiplex PCR (as described below) and API 20E gallery (Biomérieux, France).

Molecular detection of *Y. pestis* was performed using an in-house multiplex PCR, using four primer sets. The primers targeted regions of the *caf1*, *pla* and *lcrV* genes located on the pFra, pPst and pYV plasmids respectively and the *irp2* chromosomal gene [44]. The PCR products were visualized in 1% agarose gel stained with SYBR safe (Thermo Fisher Scientific, Waltham, MA, USA).

#### 2.7.2. Escherichia coli

The lung sections of 11 NPs were tested using the IHC technique with an anti-*Escherichia coli* (Rabbit Antibody to *E. coli* 1001-Virostat^®^, ME, USA) for sepsis investigation, following standardized protocols [45].

### 2.8. Protozoan Agents

#### 2.8.1. Toxoplasma gondii

The lungs and liver sections of 11 NPs were submitted to the IHC technique with an anti-*T. gondii* (Dako, Carpinteria^®^, California, USA) using standardized protocols [46]. Serum samples of seven NPs (three *A. ululata*, two *A. guariba*, one *A. caraya* and one *B. arachnoides*) were subjected to an indirect fluorescent antibody test (IFAT) in the *Laboratório de Toxoplasmose e outras Protozooses* (LabTOXO), *IOC – Fiocruz*. Following Camargo (1964), IgG anti-*T. gondii* antibodies detection was conducted using an anti-monkey IgG FITC conjugate produced in rabbit (Sigma-Aldrich®, USA). *T. gondii* RH strain tachyzoites, maintained in Swiss Webster mice, were used as antigens. The follow-up of IgG anti-*T. gondii* titers was performed only in the surviving black-and-gold howler monkey (*A. caraya*, ID 2576) in the following five weeks post clinical signs onset. A serum sample of a Colombian red howler monkey (*A. seniculus*) with detectable antibodies by MAT (1:4,096) was used as positive control [47]. In view of the positive result for NPs, an employee from CPRJ that was pregnant at the time of the outbreak was also tested for the pathogen.

Whole blood samples from the same seven animals tested in serology were used to detect *T. gondii* DNA. DNA extraction was performed using Qiamp DNA Blood Mini Kit (Qiagen^®^ Inc., USA), following the manufacturer’s instructions. *T. gondii* DNA detection was performed by conventional PCR to amplify a 529 base pairs (bp) repeat element (REP529) previously described [48]. DNA amplification was observed by electrophoresis in agarose 1% gel stained with GelRed^®^ (Biotium, USA). The DNA from *T. gondii* RH strain tachyzoites was used as positive control. The produced amplicons were purified by cycling with ExoSAP-IT enzyme (Applied Biosystems, USA). All the samples were sequenced using the same primers as those used in the PCR reactions, in a 3730xl DNA analyzer (Applied Biosystems, USA). Sequences were analyzed through Bioedit v.7.1.9 [49], and compared with the NCBI database using BLASTn tool (https://blast.ncbi.nlm.nih.gov/Blast.cgi).

## 3. Results

### 3.1. Clinical description of the CPRJ outbreak in neotropical primates

Eleven primates from six different species housed in six enclosures in two sectors of the CPRJ were affected by the outbreak from September to November 2020 (**Fig 1A**). One species is listed as Critically Endangered (*B. arachnoides*) and one is listed as Endangered (*A. ululata*) in the Red List of Threatened Species by the International Union for Conservation of Nature and Natural Resources (IUCN). One primate (female *P. caligatus*) did not show clinical signs and was found dead a few hours after normal feeding. Ten primates presented at least one clinical sign, two of them (male and female *C. melanocephalus*) have died before a complete clinical examination could be performed. Inappetence and anorexia were the first clinical signs to be observed in symptomatic animals and were observed in all of them. Prostration was also observed in all symptomatic individuals, ranging from mild (lethargy, but still moving around the enclosure – 1/10 animals; 10%), moderate (animal visibly depressed, spending most of the time motionless, but still perched – 7/10 animals; 70%) and accentuated (animal totally lethargic, unable to perch, remaining on the enclosure floor – 2/10 animals; 20%). Two primates (20%) showed drowsiness and inability to keep their eyes open.

**Fig 1.**
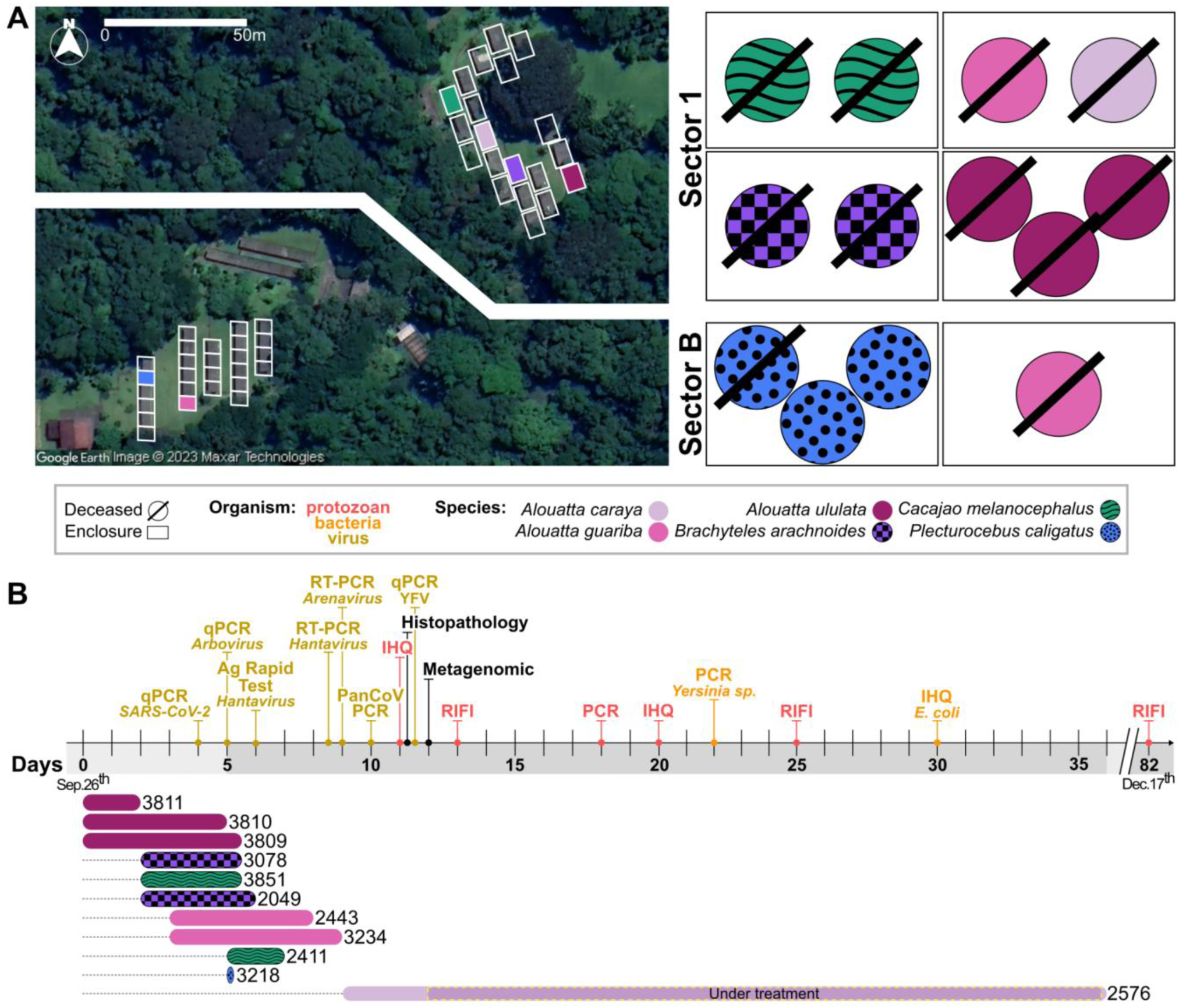
Geographical distribution of the cases and general course of the outbreak. (**A**) Spatial location of the enclosures (left) and their graphical representation (right). The rectangles represent the enclosures, which are colored according to the neotropical primate (NP) species affected by the outbreak: *Alouatta* genus in different shades of pink, *Brachyteles* in orange, *Cacajao* in green and *Plecturocebus* in blue. Unfilled rectangles had no animals affected by the outbreak. (**B**) The top-half of the timeline chronologically shows all major tests performed during the outbreak, separated by color according to the pathological agent: protozoan (*Toxoplasma gondii*) in red, bacterial agents in green and viral agents in blue. Exception is the metagenomic sequencing and histopathology, highlighted in black. Actual timeline is represented as a horizontal bar in consecutive days from the onset of symptoms in the first primate (day 0) until the death of the last animal (day 82). Double bars represent a time cut. The bottom-half of the display is dominated by the graphic representations of the symptomatic periods of each animal. The bars represent the beginning of the clinical signs until the death of the respective primate. Numbers on the right side of each bar represent the animal identification, and colors and patterns inside the bars represent the different NP genera. Only one animal (3218) died without showing any clinical sign.

Abdominal pain and distension were observed in 20% and 40% of symptomatic primates, respectively. Respiratory alterations such as dyspnea and wheezing on pulmonary auscultation were present in 20% of the symptomatic animals, while nasal secretion was observed in only one primate (10% of the symptomatic ones). Body temperature was measured in 8 of the 11 affected animals and ranged from 36.2 to 41.4 °C (97.16 F to 106.52 F). For ten of the eleven animals (91%), the time between the onset of symptoms and death ranged from zero to seven days with an average time of 3 days. One animal survived longer (36 days) and was later euthanized due to poor prognosis (for details, see [50]). The general course of the outbreak, as well as the main testing are summarized in **Fig 1B**.

### 3.2. Pathological Description of the CPRJ outbreak in neotropical primates

The main pathological findings are described in **Table 1** and were carried out by a multi-institutional team formed by members of CPRJ, SAP/UFRuralRJ and SCPrim/ICTB. The liver of the 11 NPs was the hallmark gross organ. It was markedly enlarged, with white to yellow random dots (hepatocellular necrosis) mottled with random red dots (hemorrhage) at the capsular (**Fig 2A**) and cut surface. In the lungs, the pleural surface was irregularly covered by multifocal red areas (hemorrhage) (**Fig 2B**). White multifocal dots were seen at the spleen’s capsular, and parenchyma cut surface (**Fig 2C**). Mediastinal and mesenteric lymph nodes were enlarged, and the cut surface contained multifocal white and red areas (**Fig 2D**). The stomach showed marked focal ulcerative gastritis (**Fig 2E**). The jejunum showed petechial to ecchymotic multifocal areas at the serosal surface (**Fig 2F**). With the presented scenario, an epizootic notification was made to the State Department of Health of Rio de Janeiro, as officially required.

**Fig 2.**
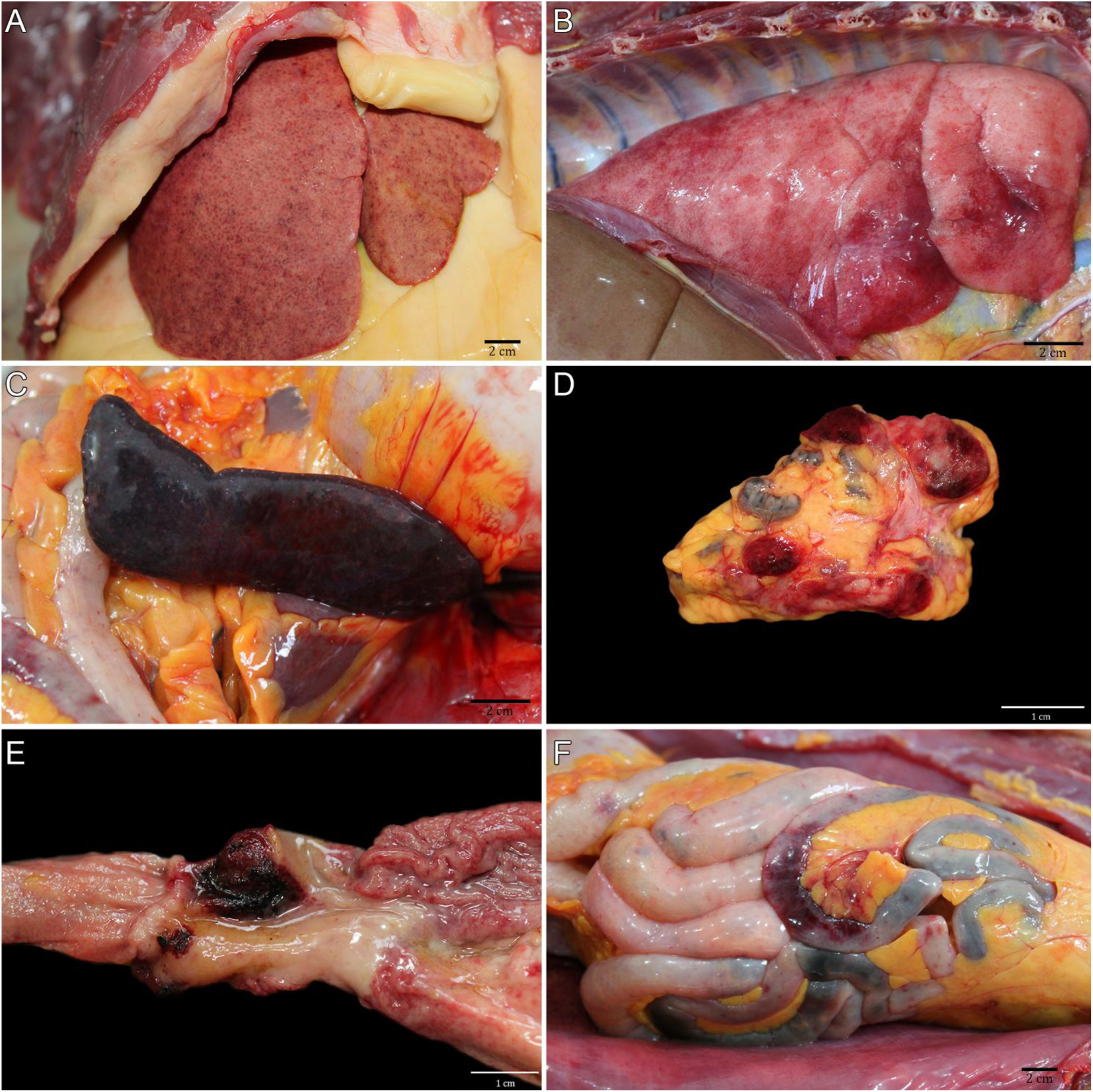
Gross findings of toxoplasmosis in Neotropical Primates (NP). (A) NP 2. Liver diffusely enlarged, with random hepatocellular necrosis and hemorrhage areas. (B) NP 5. Multifocal hemorrhagic areas on pleural surface. (C) NP 11. Diffuse splenomegaly with white multifocal dots at the capsular surface. (D) NP 11. Mesenteric lymph node enlarged with multifocal white and red areas at the cut surface. (E) NP 11. Evident focal ulcer in the pylorus. (F) NP 11. Petechial to ecchymotic multifocal areas at the jejunum serosal surface.

**Table 1.** Main pathological findings in Neotropical Nonhuman Primates with toxoplasmosis.

| NHP | Species | Sex | Age* | Body condition score** | Hepatomegaly | Splenomegaly | Lymphadenomegaly | Hemorrhages |  |  |  |  | Necrosis |  |
| --- | --- | --- | --- | --- | --- | --- | --- | --- | --- | --- | --- | --- | --- | --- |
|  |  |  |  |  |  |  |  | Lungs | Liver | Lymph node | Small intestine | Large intestine | Stomach | Intestine |
| Atelidae | 1 *B. arachnoides* | M | 6 | - | + | + | + | + | + | + | + | + | - | + |
|  | 2 *B. arachnoides* | F | 20 | 4 | + | + | + | + | + | + | + | - | - | + |
|  | 3 *A. ululata* | F | Adult | - | + | + | + | + | - | + | + | + | - | - |
|  | 4 *A. ululata* | F | Adult | - | + | + | + | - | - | + | - | - | - | - |
|  | 5 *A. ululata* | F | Adult | 3 | + | + | + | + | + | + | - | - | - | - |
|  | 6 *A. guariba* | M | 11 | 3 | - | + | - | - | - | - | - | - | - | - |
|  | 7 *A. guariba* | M | 7 | 3 | - | - | - | - | - | - | - | - | - | - |
|  | 8 *A. caraya* | M | 10 | 2 | - | + | + | - | + | + | - | - | - | - |
| Pitheciidae | 9 *C. melanocephalus* | F | 12 | 2 | - | + | + | - | - | + | - | - | - | - |
|  | 10 *C. melanocephalus* | M | Adult | - |  |  |  | + | - | - | - | - | - | - |
|  | 11 *P. caligatus* | F | 4 | 3 | + | + | + | + | + | + | + | - | + | + |

(M) Male; (F) Female; \*Years; (-) Absent; (+) Present. \*\* 1 to 5 scale

### 3.3. Negative SARS-CoV-2 and arboviral diagnoses

As the first affected NPs had clinical respiratory signs and considering the COVID-19 pandemic scenario in Brazil, we first conducted SARS-CoV-2 testing in three laboratories from UFRJ (LDDV, LVM and NEEDIER). We obtained negative molecular diagnostic results for primates and CPRJ workers. Serological results were also negative for all NPs, but positive for two animal keepers, consistent with previously undiagnosed SARS-CoV-2 infection. Additional molecular testing for universal diagnosis of coronavirus was also performed on swab and tissue samples from primates, again with negative results (**Fig 1B**).

In view of the pathological findings, especially the intestinal hemorrhages, a multiplex qPCR test for seven arboviruses (YFV, dengue, Zika, chikungunya, Mayaro, Oropouche and West Nile) was performed, all with negative results, which was confirmed by the Regional Reference Laboratory for Yellow Fever of IOC. Following investigation, NP blood samples were forwarded to the reference *LHR*/*IOC – Fiocruz* for hantavirus and arenavirus investigation, including lymphocytic choriomeningitis virus, which were all negative.

Concomitantly, recommendations regarding biosafety issues were made to ensure the safety of CPRJ workers and NPs, as well as the scheduling of clinical care and collection of blood samples for additional analysis of all professionals with a clinical condition or with a history of contact with animals. A 10-week pregnant worker had no clinical manifestation, while four other workers had mild influenza-like clinical symptoms. Serological tests for toxoplasmosis and hepatitis A, B and C were all non-reactive.

### 3.4. Histological findings for neotropical primates

Histological examinations were performed by SAP/UFRuralRJ and findings are described in **Table 2**. Extracellular structures, rounded to fusiform, basophilic, with 2 to 3 µm (tachyzoites) and basophilic, thin walls oval structures, with an average of 20 x 15 µm filled with elongated basophilic bradyzoites ranging in size from 1 to 2 µm (bradyzoite cysts) were visualized. In the liver, there was multifocal random hepatocellular necrosis and lymphocytic periportal hepatitis with intralesional tachyzoites (**Fig 3A**) and cysts of bradyzoites (**Fig 3B**), marked multifocal lipidosis, and multifocal hemosiderosis. Necrotizing fibrinoid vasculitis was observed in the liver of two primates. In the lung, there was multifocal thickening of alveolar septa by lymphocyte infiltration, intra-alveolar foamy macrophages, alveolar edema and hemorrhage, necrosis of type I pneumocyte and fibrin deposition in the alveolar space, occasionally forming hyaline membrane. Intralesional cysts of bradyzoites and tachyzoites were seen (**Fig 3C**). Necrotizing splenitis with neutrophilic and histiocytic infiltration was seen in the red and white pulp, fibrin deposition, and intralesional cysts of bradyzoites and tachyzoites (**Fig 3D**). The lymph nodes showed medullary to cortical necrosis, neutrophil, and histiocyte infiltration, mainly at the subcapsular sinus, with intralesional cysts of bradyzoites and tachyzoites. Necrohemorrhagic duodenitis with intralesional cysts of bradyzoites and tachyzoites (**Fig 3E**) was seen in three primates. Three primates presented brain lesions characterized by multifocal areas (telencephalon gray and white matter, corpus callosum, cerebellum molecular layer and myelencephalon) of malacia with intralesional tachyzoites (**Fig 3F**). In one case, the neuronal lesions were characterized by multifocal gliosis at telencephalon gray and white matter. Necrohemorrhagic tiflitis with intralesional cysts of bradyzoites and tachyzoites were seen in one primate. Another primate showed lymphocytic interstitial nephritis.

**Fig 3.**
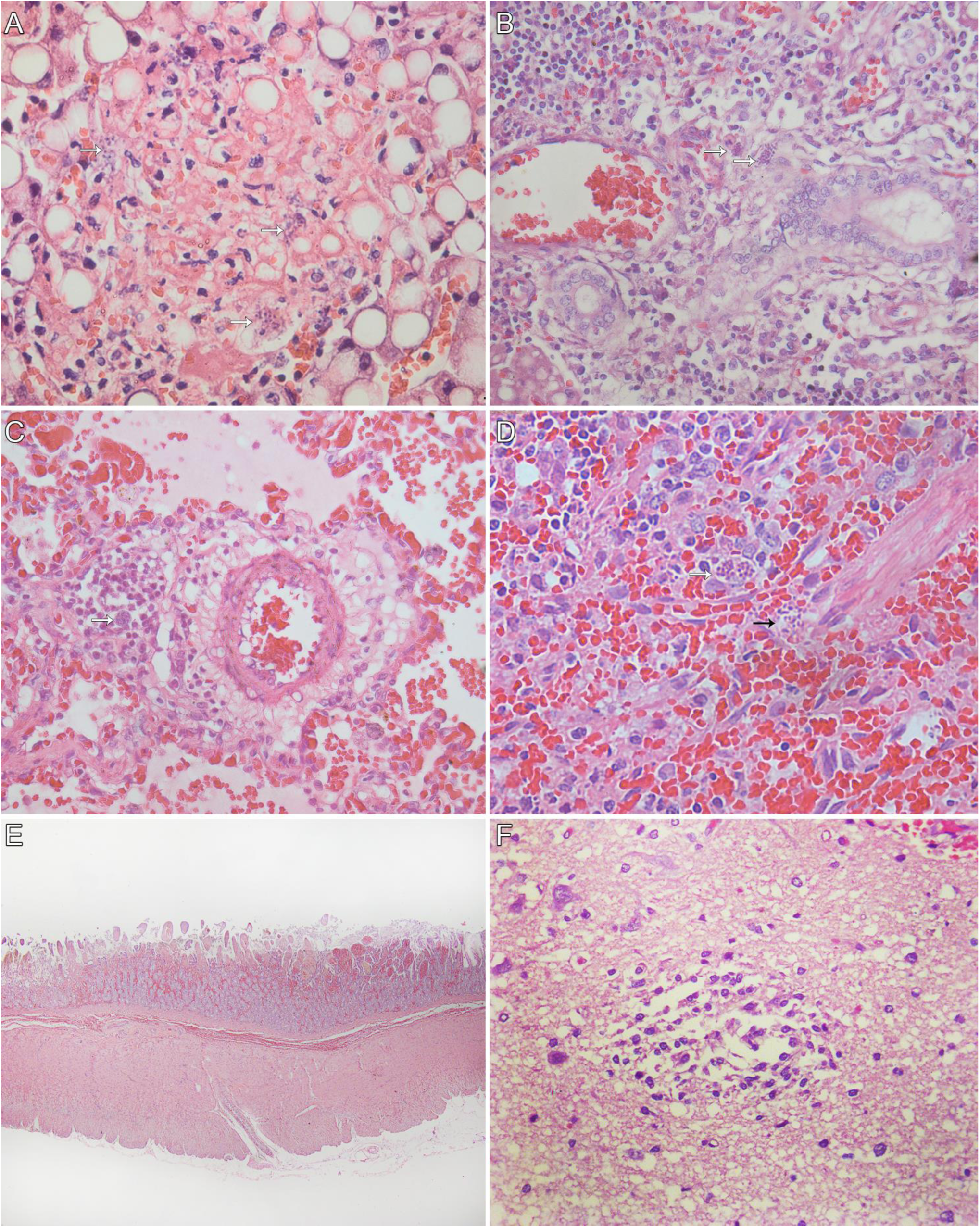
Histological findings of toxoplasmosis in Neotropical Primates (NP). (A) NP 5. Marked hepatocellular necrosis with intralesional tachyzoites (HE, Obj. 40x). (B) NP 5. Lymphocytic periportal hepatitis with intralesional cysts of bradyzoites (white arrow) (HE, Obj. 40x). (C) NP 11. Alveolar septa expanded by lymphocytes and intralesional cysts of bradyzoites (white arrow). There are multifocal areas of edema and hemorrhage in the adjacent airways. (HE, Obj. 40x) (D) NP 2. Necrotizing splenitis with intralesional cysts of bradyzoites (white arrow) and tachyzoites (black arrow) (HE, Obj. 40x). (E) NP 11. Duodenal mucosa and submucosa expanded by marked hemorrhage (HE, Obj. 2,5x). (F) NP 9. Focal malacia area at the telencephalic cortex gray matter. (HE, Obj. 40x).

**Table 2.**
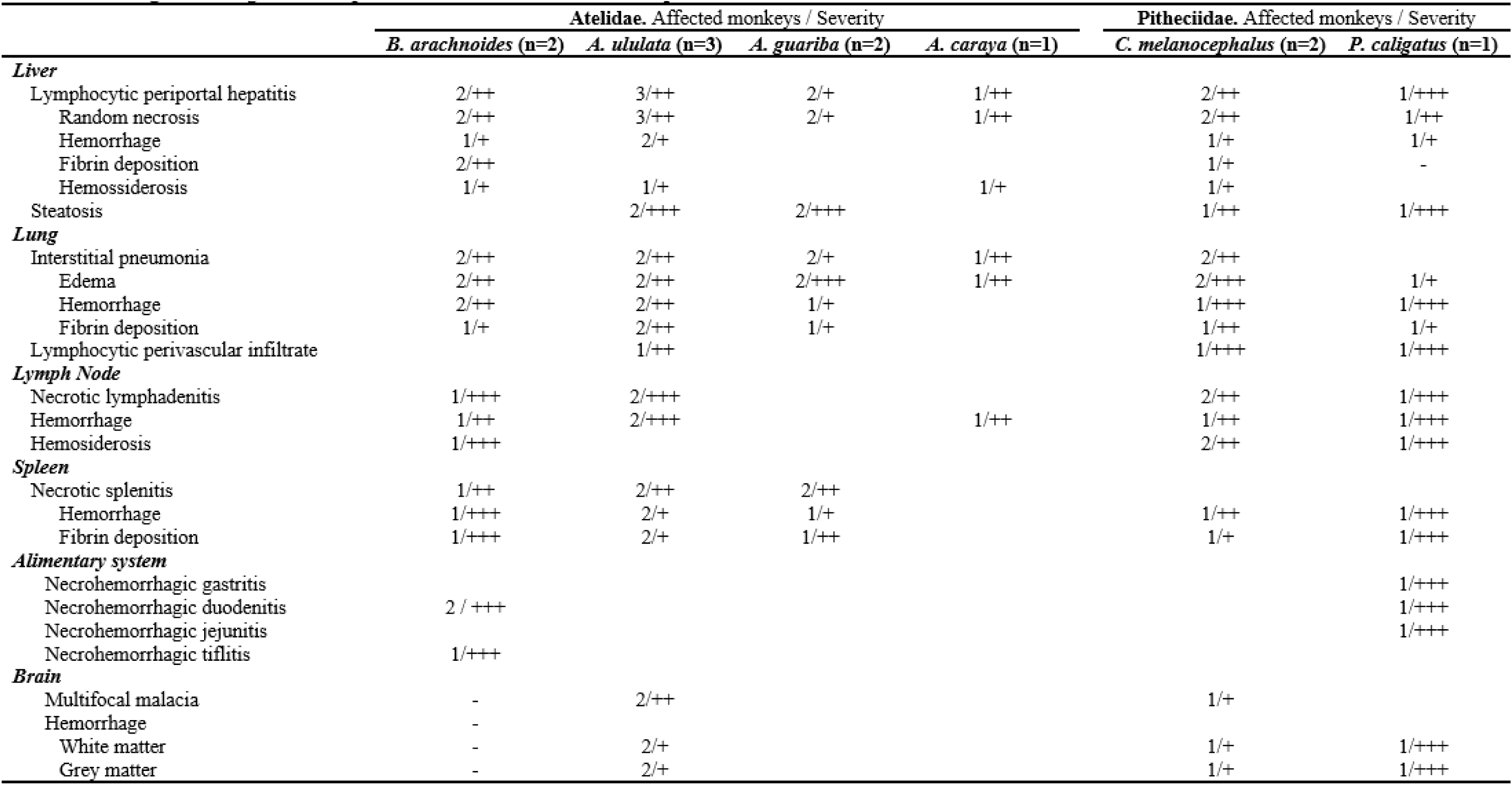
Histological findings in Neotropical Nonhuman Primates with toxoplasmosis.

### 3.5. Pathogen metagenomics reveals sepsis causing bacterial infection

In light of the emergency for a diagnosis and the need for a more detailed investigation, untargeted metagenomic sequencing was performed in a partnership between LDDV/UFRJ and IMT/USP. All samples analyzed presented a high proportion of unclassified reads, corresponding to approximately 67% (min. 58% and max. 85%) of the total reads generated (**S2 Fig**). The remaining reads, in most libraries (n=13/20), were identified as eukaryotes (*Homo sapiens)*, followed by bacteria and viruses. Seven libraries (n=7/20) showed a higher proportion of bacterial reads, representing up to 26% of all reads generated in each library. For these samples, an inversion in taxonomic representativeness was observed, with bacteria being the most represented taxonomic level, followed by eukaryotes and viruses.

Regarding the identified bacteria, diversity varied considerably between samples, without an overall dominant order or family. Nonetheless, the presence of some bacteria caught our attention, such as *E. coli* and the *Yersinia* genus. Although *E. coli* is naturally found in the intestine, it is not usually found in other organs. It is important to point out that more than half of these libraries (57%; 4/7) were generated from plasma samples, a tissue that, in theory, should be sterile. This observation, together with the necropsy findings, led us to conclude that these animals suffered from a bacterial infection that caused sepsis.

Following the histopathology observation of different *T. gondii* forms, we performed a reference assembly with the metagenomic data (reference accession number: NC_031467.1), which confirmed the presence of this pathogen. We found over 660,000 reads across all samples (median number of mapped reads: 33,000) and up to 109,870 reads in just one liver sample (from 2.5 to 17 times more than other tissue samples) (**S1 Table**).

### 3.6. Immunohistochemistry

The immunohistochemistry was performed by SAP/UFRuralRJ and findings are described in **Table 3**. Intralesional cysts of bradyzoites and tachyzoites were visualized in more than one organ of almost all primates that died naturally (10/11). In only one case of *A. caraya* with subacute evolution, *T. gondii* structures were not seen. The clinical-pathological findings were already reported [50]. Infection by SARS-CoV-2 by IHQ was ruled out in all 11 tested NP. Out of eleven primates, seven were tested for *E. coli* infection and confirmed for coinfection of *T. gondii* and *E. coli* pneumonia, as well as sepsis, which was confirmed by vessels intraluminal *E. coli* immunolabelling and contributed to the worsening of their health conditions.

**Table 3.**
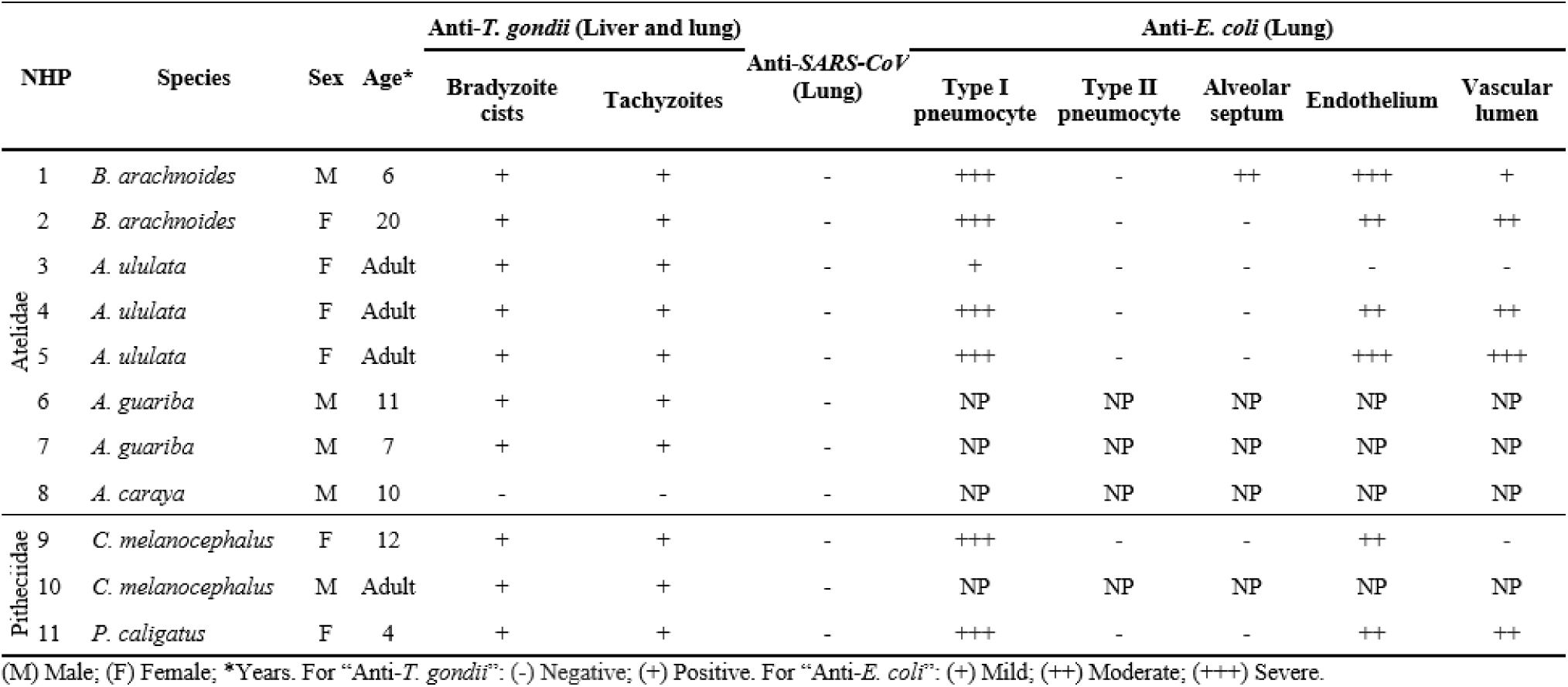
Immunohistological findings in Neotropical Nonhuman Primates with toxoplasmosis.

| NHP | Species | Sex | Age* | Anti-*T. gondii* (Liver and lung) |  | Anti-SARS-CoV<br>(Lung) | Anti-*E. coli* (Lung) |  |  |  |  |
| --- | --- | --- | --- | --- | --- | --- | --- | --- | --- | --- | --- |
|  |  |  |  | Bradyzoite<br>cysts | Tachyzoites |  | Type I<br>pneumocyte | Type II<br>pneumocyte | Alveolar<br>septum | Endothelium | Vascular<br>lumen |
| 1 | *B. arachnoides* | M | 6 | + | + | - | +++ | - | ++ | +++ | + |
| 2 | *B. arachnoides* | F | 20 | + | + | - | +++ | - | - | ++ | ++ |
| 3 | *A. ululata* | F | Adult | + | + | - | + | - | - | - | - |
| 4 | *A. ululata* | F | Adult | + | + | - | +++ | - | - | ++ | ++ |
| 5 | *A. ululata* | F | Adult | + | + | - | +++ | - | - | +++ | +++ |
| 6 | *A. guariba* | M | 11 | + | + | - | NP | NP | NP | NP | NP |
| 7 | *A. guariba* | M | 7 | + | + | - | NP | NP | NP | NP | NP |
| 8 | *A. caraya* | M | 10 | - | - | - | NP | NP | NP | NP | NP |
| 9 | *C. melanocephalus* | F | 12 | + | + | - | +++ | - | - | ++ | - |
| 10 | *C. melanocephalus* | M | Adult | + | + | - | NP | NP | NP | NP | NP |
| 11 | *P. caligatus* | F | 4 | + | + | - | +++ | - | - | ++ | ++ |

(M) Male; (F) Female; \*Years. For “Anti-*T. gondii*”: (-) Negative; (+) Positive. For “Anti-*E. coli*”: (+) Mild; (++) Moderate; (+++) Severe.

### 3.7. Molecular and serological diagnosis of toxoplasmosis

In view of the positive result for toxoplasmosis, blood samples from all animals were also tested molecularly in the LabTOXO/*IOC – Fiocruz*. From the seven serum samples of NP initially submitted to IFAT, only the sample from a black-and-gold howler monkey (*A. caraya*, ID 2576) showed IgG anti-*T. gondii* (1/7, 14.3%), with titers of 1:16. Serological follow-up of this animal showed a progressive increase in antibody titers in the five consecutive weeks, reaching titers of 1:64 and 1:256 in the second and fourth weeks of illness, respectively. Additionally, the 529 bp repeat element of *T. gondii* DNA was detected in all whole blood samples submitted to PCR. Of these, six showed high identities with *T. gondii,* ranging between 98% and 100% when compared with nucleotide sequences deposited in GenBank (LC547467.1). Therefore, with the clinical-pathological presented picture and the multiple auxiliary examination for *T. gondii* detection, we concluded that toxoplasmosis was the most likely etiological agent causing this outbreak.

### 3.8. Negative *Yersinia* bacteriological, molecular, and serological diagnosis

Regarding the *Yersinia* genus, we found metagenomic reads from different species in all samples submitted to metagenomics. A reference-guided assembly was conducted with *Y. pestis* (AE017042.1), *Y. enterocolitica* (NC_008800.1) and *Y. pseudotuberculosis* (NZ_LR134373.1) complete genomes, some of the main yersiniae of zoonotic importance. We found that *Y. pestis* recovered the least amount of reads in all samples (approximately 228,000 reads total, max. 40.532 and min. 1.229). On the other hand, the assembly using *Y. enterocolitica* genome as reference was the most effective, with over 723,000 reads total and up to 140,637 reads recovered in just one sample (**S1 Table**). Because of these observations and the CPRJ’s proximity to a previous plague focus, we could not rule out a *Y. pestis* infection at this point. Official guidelines support that to confirm plague, it is necessary to have a positive culture or positive results in both serological and molecular tests. In this manner, no *Y. pestis* growth was observed and all samples showed no seroreactivity in the hemagglutination/ELISA tests performed by the IAM – *Fiocruz* and no amplification was observed for *Y. pestis* specific genes *pla* and *caf1*. These results, associated with the higher prevalence of *Y. enterocolitica*, suggest that this pathogen was not the main source of this outbreak.

## 4. Discussion

Due to continuous human population expansion, enhanced mobility and intense land usage patterns, pathogen spillover has become increasingly frequent in recent decades, often leading to an increase in zoonotic and epizootic outbreaks [51]. Zoonotic disease surveillance targets primarily mammals and especially NHPs, which harbor a wide variety of pathogens [52]. In this study, we present an effort to create an integrated epidemiological and genomic investigation group to solve an NP outbreak in the CPRJ / Brazil, during the first year of the COVID-19 pandemic. The workgroup was able to mitigate the health alert and reach an accurate diagnosis by reporting a fulminant case of *T. gondii*.

In order to identify the causative agent of this outbreak, we first suspected a SARS-CoV-2 infection, considering the pandemic scenario in which the outbreak happened (during the emergence of the new variant P.2 in Brazil) and the fact that two CPRJ staff members had reported going to work with respiratory symptoms, which was soon discarded. Regarding the main two compulsory notifiable diseases, rabies has a neurologic clinical picture, such as behavioral changes, salivation and paralysis [5], that was not seen in any of the affected primates. However, YFV is one of the major causes of death in NPs and presents general clinical signs, such as prostration, loss of appetite, dehydration, diarrhea and jaundice [5]. Many of these were observed in the CPRJ NPs, justifying the investigation using a suite of diagnostic techniques. Since other arboviruses can also affect NPs [53–55], we also tested them for dengue, Zika, chikungunya, Mayaro, Oropouche and West Nile. Infections with *Yersiniae*, hantavirus and arenavirus were also investigated, considering the intestinal hemorrhage detected and CPRJ’s proximity to urban centers and forests, where there is a large circulation of wild rodents. All these suspicions were dismissed.

In contrast, *T. gondii* DNA was detected in whole blood samples from all specimens evaluated. This same type of biological sample was also used for the detection of parasitic DNA in a case of fatal toxoplasmosis in a free-living southern muriqui in São Paulo/Brazil [56]. The detection of *T. gondii* DNA in blood samples can be a strong indication of the acute phase of the parasitosis in recently infected hosts [57]. These results highlight the potential use of whole blood PCR as a diagnostic tool in cases of acute toxoplasmosis in captive NP, allowing the immediate start of specific therapeutic management against the parasite in symptomatic animals, which may increase the survival of individuals.

Primates are kept in familiar groups and housed in outdoor enclosures built in mesh with a natural floor covered with leaves, where they eventually encounter other sylvatic animals such as birds, snakes, bats and rodents, besides mosquitoes and other insects. However, there were no reports of felines circulating in the center or in its surroundings, so an infection with *T. gondii* was not immediately suspected. Several cases of toxoplasmosis in NHP have already been reported and reviewed around the world [58–60]. It is a cosmopolitan infectious disease that causes either an acute or chronic clinical manifestation and affects a wide variety of mammals and birds [59]. Among wild animals, toxoplasmosis is particularly dangerous for NP, since almost all cases reported to date have been acute and fatal, with nonspecific signs (mainly apathy, anorexia, abdominal distension, and fever), making diagnosis challenging [61–66]. With the confirmation of the toxoplasmosis, it is believed that infection may have occurred through contaminated foliage, consumed by this group of animals as part of their regular diet. This foliage is purchased at local markets, where there is a large circulation of feral and stray cats and may not have been properly sanitized before consumption. This could explain why the affected animals were distributed in different enclosures, far apart from each other, and why the disease did not spread to other enclosures. However, by the time when toxoplasmosis was detected, there was no foliage left to be analyzed. Furthermore, it is important to note that, as discussed by Amendoeira *et al.*, the strain of *T. gondii* isolated from the tissues of specimen 2576 (*A. caraya*), which inoculum in mice proved to be highly virulent [67], is genetically similar to the ones found in seven asymptomatic birds in Espírito Santo (ES) [68–70] and four human congenital toxoplasmosis cases in Minas Gerais (MG) in 2012 [70,71]. Both ES and MG states border Rio de Janeiro state, where the CPRJ is located.

Due to local biosecurity actions and the joint work of different research and health institutions, we identified toxoplasmosis in 2 days after necropsies. It is important to note that because of the security measures adopted due to the pandemic, the center had a significant reduction in the number of working employees, making all procedures take longer than usual, such as necropsies and histopathology. Although ten animals died, the black-and-gold howler monkey (2576; *A. caraya*) began treatment just two days after the onset of clinical signs, being the first report of a NP surviving from toxoplasmosis [50]. This was the only animal who showed IgG antibodies in the initial serological screening, progressively increasing antibody titers over the following five weeks [50]. However, there was no recovery of the general health status of the animal after implementation of treatment, resulting in the euthanasia of the individual. It is possible that factors such as the difference in parasite load to which the individual was exposed in the captivity context, added to the rapid administration of specific treatment for toxoplasmosis, contributed to the survival of this specimen, which resulted in the detection of IgG and increase in antibody titers in the individual’s serum samples. In the other NP evaluated, the acute profile with fatal outcome of toxoplasmosis may have prevented the individuals from having enough time to start producing antibodies, which is reflected in serology negative results. Another study also failed to detect anti-*T. gondii* antibodies in a captive *Callicebus nigrifons* (Pitheciidae) that died suddenly of toxoplasmosis in Brazil [72]. This report, together with the serological results of the present study, confirm the absence of a humoral response in some NP species during acute *T. gondii* infection. This fact becomes an important obstacle for the serodiagnosis of toxoplasmosis in these species in captivity.

While a national epizootic surveillance strategy started in mid-1999 in Brazil, it was only two decades ago that the Ministry officially recognized animals as sentinel organisms, with importance for the epidemiological surveillance system [2,73]. Although the recognition itself was already a great advance, it is a short period for major progress. This partially explains why the workflow is still limited in practice. Testing majorly for YFV and rabies is one of the many bottlenecks for epizootic diagnosis. For very few cases, samples that cross this first barrier are very often directed for herpesvirus detection, as it is a very characteristic disease that can have some overlapping symptoms related to YFV, such as vomiting and prostration [74]. Herpesviruses infection leads to a chronic and frequently asymptomatic illness in humans, yet it is usually fatal for NHPs. In this sense, all mild symptoms and hidden diseases are largely overlooked, reaching another bottleneck for effective epizootic surveillance.

Strictly speaking, only cases or suspicions of diseases that appear on the National List of Compulsory Notification should be reported, which excludes a great number of other conditions. In practice, all NHP cases have to be reported due to the risk of YFV. Yet, even for notifiable diseases, data are not widely and clearly available, making access to information difficult. Ideally, as we have a large number of agents that can affect animal populations, all notified cases should go through a wide investigation for multiple pathogens, which is typically not done. It is very rare for the community to perform those screenings, both because of the lack of appropriate funding for such high-cost experiments and the lack of trained staff or lab expertise. Importantly, considering that more than 24% of the NP are considered endangered or worse in the Red List of Threatened Species by the IUCN [75], their conservation should be a key concern.

Therefore, diseases that threaten animal survival, along with those that pose risks to humans should both be closely monitored. A more integrative surveillance system would facilitate interactions between reference laboratories and research institutions across the country, enabling joint action in more complex cases or those that do not fit into the hall of traditionally investigated diseases. High-resolution techniques are still expensive and do not replace the traditional ones, but could benefit in these cases. Decisions about pharmaceutical or non-pharmaceutical interventions could also benefit from this integration and be made collectively, contributing to outbreak control in non-human populations. It is essential to point out that this outbreak was quickly solved due to the actions of researchers that invested their time and resources during a sensitive time, such as the quarantine. Thus, an immediate collective and interdisciplinary response was essential to contain this outbreak, preventing its further spread to other animals, and the rapid identification of the causative agent allowed the implementation of innovative treatments that contributed to ameliorate the clinical outcome for the last affected animal.

## 5. Conclusions

This investigation concluded that the outbreak was caused by *T. gondii* after 48h after necropsy procedures and is the first description of toxoplasmosis coinfection with bacterial sepsis. We emphasize that gross examination and histopathology should be the starting choice for a diagnosis flowchart, while IHQ and PCR should be employed for the quick etiological confirmation. Diagnosis for toxoplasmosis is not included in the hall of zoonotic diseases investigated under official guidelines. Our study showcases a cross-platform interdisciplinary investigation to detect pathogens with public health relevance that are not included in current diagnostic policies, suggesting that the ongoing model of testing mainly for YFV and rabies presents important flaws and could be improved. Current public health policies on non-human animal disease notifications could make full use of existing interdisciplinary outbreak investigation approaches within a One Health framework.

## 6. Data sharing

Raw sequencing data generated in the study was submitted to the Sequence Read Archive (SRA) databank from the NCBI server under the BioProject number PRJNA986486 and BioSamples numbers SAMN35839772 to SAMN35839791. Other relevant data are within the manuscript or available at https://github.com/lddv-ufrj/CPRJ_Outbreak.

## 7. Funding

This work was supported by the *Conselho Nacional de Desenvolvimento Científico e Tecnológico*/CNPq (grant number 313005/2020-6 to A.F.A.S.); *Fundação de Amparo à Pesquisa do Estado do Rio de Janeiro*/FAPERJ (grants numbers E26/211.040/2019, E-26/210.245/2020 and E-26/201.193/2022 to A.F.A.S and E-26/204.136/2022 to F.B.S.); the Medical Research Council-São Paulo Research Foundation (FAPESP) CADDE partnership award (MR/S0195/1 and FAPESP18/143890 to N.R.F. and E.C.S.); Wellcome Trust and Royal Society (Sir Henry Dale Fellowship 204311/Z/16/Z to N.R.F.), and Bill & Melinda Gates Foundation (INV-034540 and INV-034652 to N.R.F.). F.B.S. was further supported by *Coordenação de Aperfeiçoamento de Pessoal de Nível Superior – Brasil* (CAPES) – Finance Code 001 (grant number 88887.507719/2020-00). The funders had no role in study design, data collection and analysis, decision to publish, or preparation of the manuscript.

## 8. Acknowledgments

We would like to give special thanks to the CPRJ staff member for their efforts in solving the outbreak and for their willingness to come to NEEDIER to collect samples. We are also grateful to the *Fundação Oswaldo Cruz*, in special the *Instituto Oswaldo Cruz*, for the support with biosafety supplies to the CPRJ staff in order to manage the outbreak and for the transport of the biological samples to the *Serviço de Referência Nacional em Peste* at *Instituto Aggeu Magalhães*. We also are grateful to Dr. Alice Laschuk Herlinger for providing primers and conditions for multiplex PCR of arboviruses. We acknowledge the support from Dr. Marcelo Alves Soares and Dr. Miguel Angelo Martins Moreira for access to the Sanger sequencing platform from *Instituto Nacional de Câncer José Alencar Gomes da Silva* (INCA). This study is part of the thesis of F.B.S. for his PhD degree from the Graduate Program in Genetics, UFRJ.

## 9. Authors’ contributions

**F.B.S.**: Formal analysis, Data curation, Investigation, Visualization, Writing – Original Draft, Writing – review & editing. **S.B.M.**: Conceptualization, Investigation, Resources, Data Curation, Writing – Original Draft, Writing – Review & Editing, Project administration. **A.H.B.P.**: Investigation, Writing – Original Draft, Writing – review & editing. **I.F.A.**: Investigation, Writing – Original Draft. **F.R.R.M.**: Formal analysis, Investigation, Data Curation, Writing – Review & Editing. **M.D.**: Formal analysis, Investigation, Writing – Review & Editing, Supervision. **I.M.C.**: Methodology, Investigation, Writing – Review & Editing. **T.A.P.**: Investigation, Resources, Project administration. **L.T.F.C.**: Investigation. **T.S.M.**: Investigation, Writing – Review & Editing. **M.A.C.C.**: Investigation, Writing – Review & Editing. **R.C.O.**: Investigation. **J.F.**: Investigation, Writing – Original Draft. **M.R.S.A.**: Investigation. **J.G.O.**: Investigation. **T.A.C.S.**: Investigation. **R.M.G.**: Resources, Supervision. **D.S.F.**: Investigation. **J.G.J.**: Investigation. **M.S.B.S.**: Investigation. **M.F.B.**: Investigation. **O.C.F.J.**: Resources, Investigation, Supervision, Data Curation, Funding acquisition. **A.T.**: Resources, Supervision, Funding acquisition. **T.M.C.**: Resources, Supervision, Writing – review & editing, Funding acquisition. **R.S.A.**: Resources. **N.R.F.**: Resources, Writing – Review & Editing, Funding acquisition. **A.M.P.A.**: Investigation, Resources, Writing – Original Draft, Writing – Review & Editing, Supervision. **A.P.**: Resources, Supervision. **E.C.S.**: Resources, Funding acquisition. **M.R.R.A.**: Investigation, Resources, Writing – Original Draft, Writing – Review & Editing, Supervision. **E.R.S.L.**: Conceptualization, Investigation, Resources, Writing – Review & Editing, Project administration, Supervision. **D.G.U.**: Investigation, Resources, Supervision, Writing – Original Draft, Writing – review & editing. **A.F.A.S.**: Conceptualization, Resources, Writing – Original Draft, Writing – Review & Editing, Supervision, Project administration, Funding acquisition.

## 10. Declarations of competing interest

The authors have declared that no competing interests exist.

## 12. Supporting information

**S1 Figure. Location of the *Centro de Primatologia*.** Brazilian map with the Rio de Janeiro state marked in a black box. The map of the Rio de Janeiro state with all municipalities colored in dark gray, except for Guapimirim, in black. The right panel is zoom in from the Guapimirim northeast border, where the CPRJ is located. Environmental protection areas are colored in green, urban areas in blue and CPRJ is represented as a yellow star.

**S2 Figure. Taxonomic diversity in NP samples classified by Kraken.** Sequences from NP samples collected during the outbreak were classified using Kraken with the miniKraken_v2 database. Eukaryotic reads are colored in light gray, bacterial reads in dark grey and viral reads in black.

**S1 Table. Main findings from the reference assembly using metagenomic data.**

**S1 Figure.**
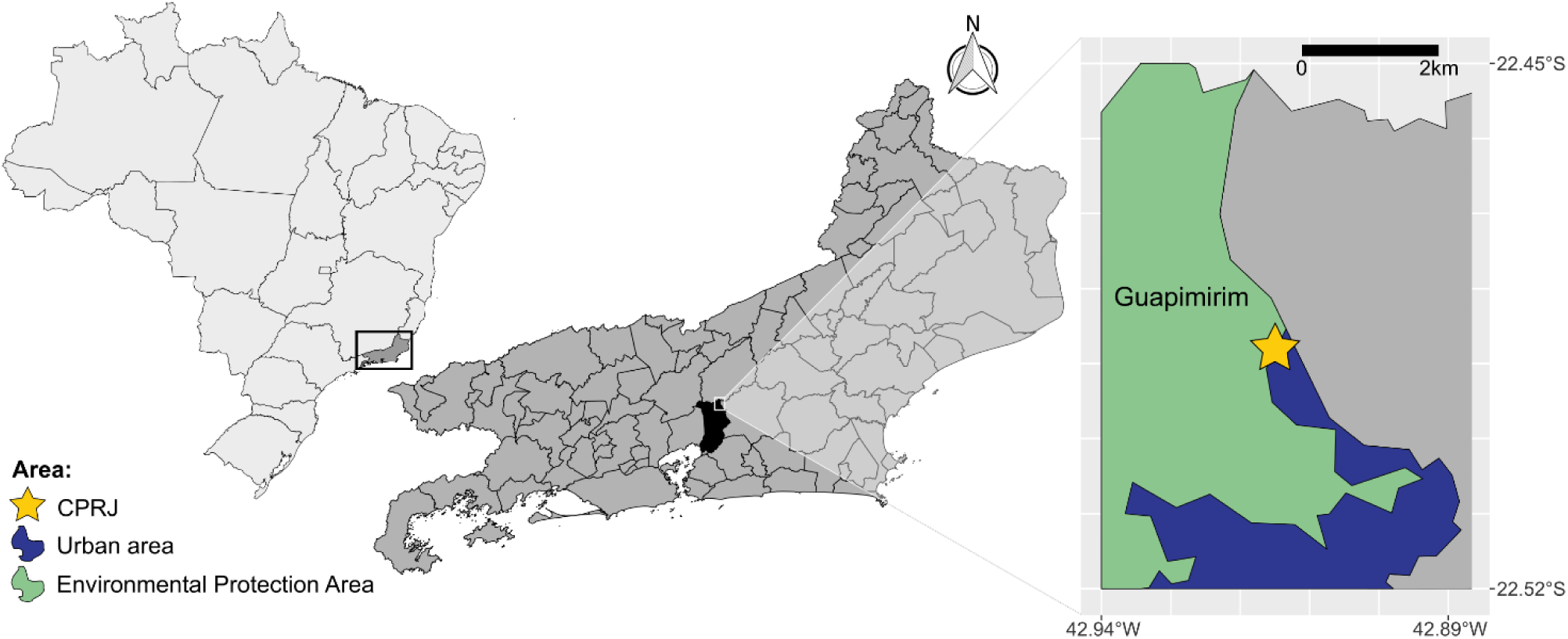
Location of the *Centro de Primatologia*. Brazilian map with the Rio de Janeiro state marked in a black box. The map of the Rio de Janeiro state with all municipalities colored in dark gray, except for Guapimirim, in black. The right panel is zoom in from the Guapimirim northeast border, where the CPRJ is located. Environmental protection areas are colored in green, urban areas in blue and CPRJ is represented as a yellow star.

**S2 Figure.**
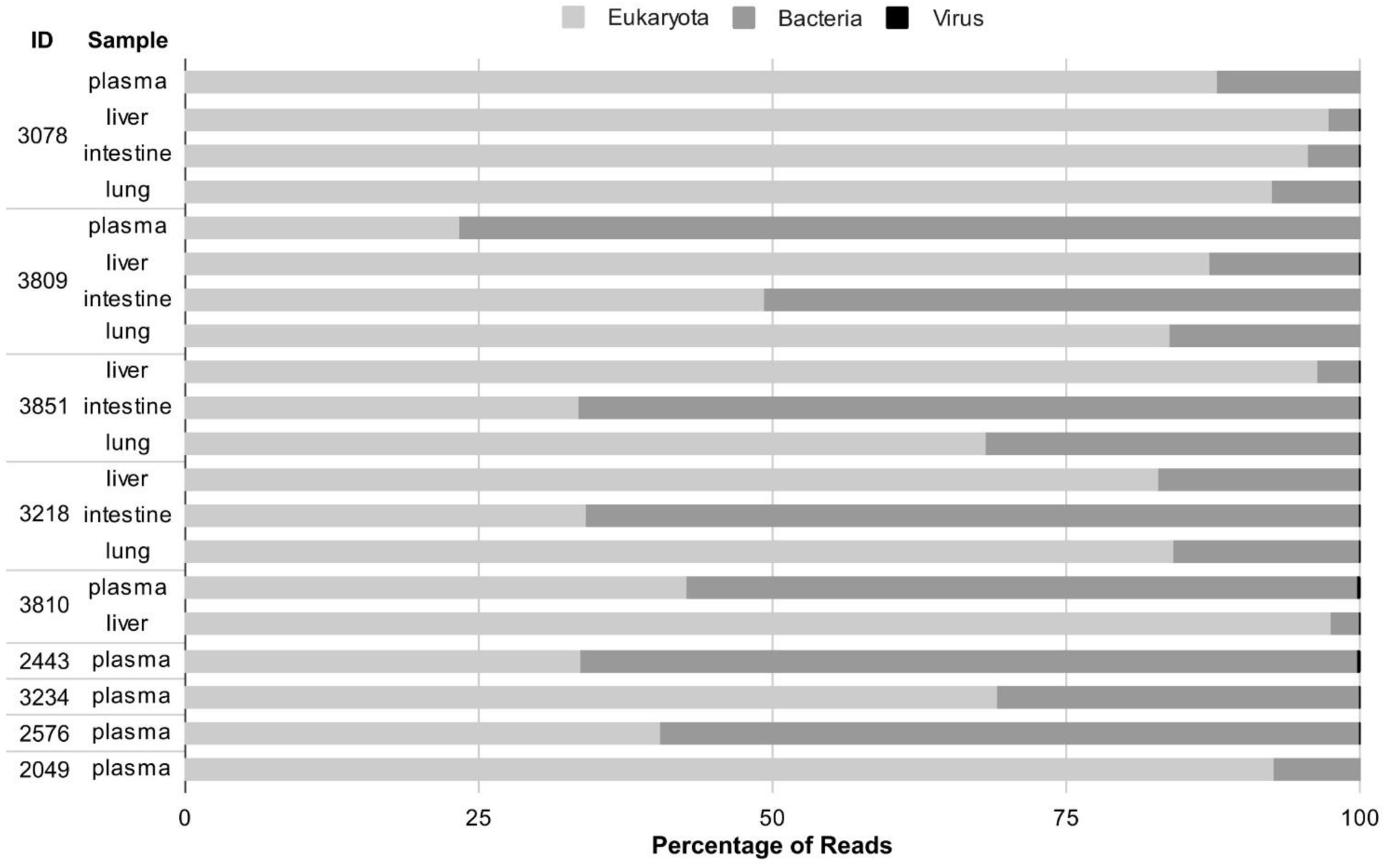
Taxonomic diversity in NP samples classified by Kraken. Sequences from NP samples collected during the outbreak were classified using Kraken with the miniKraken_v2 database. Eukaryotic reads are colored in light gray, bacterial reads in dark grey and viral reads in black.

**Supplementary Table 1.**
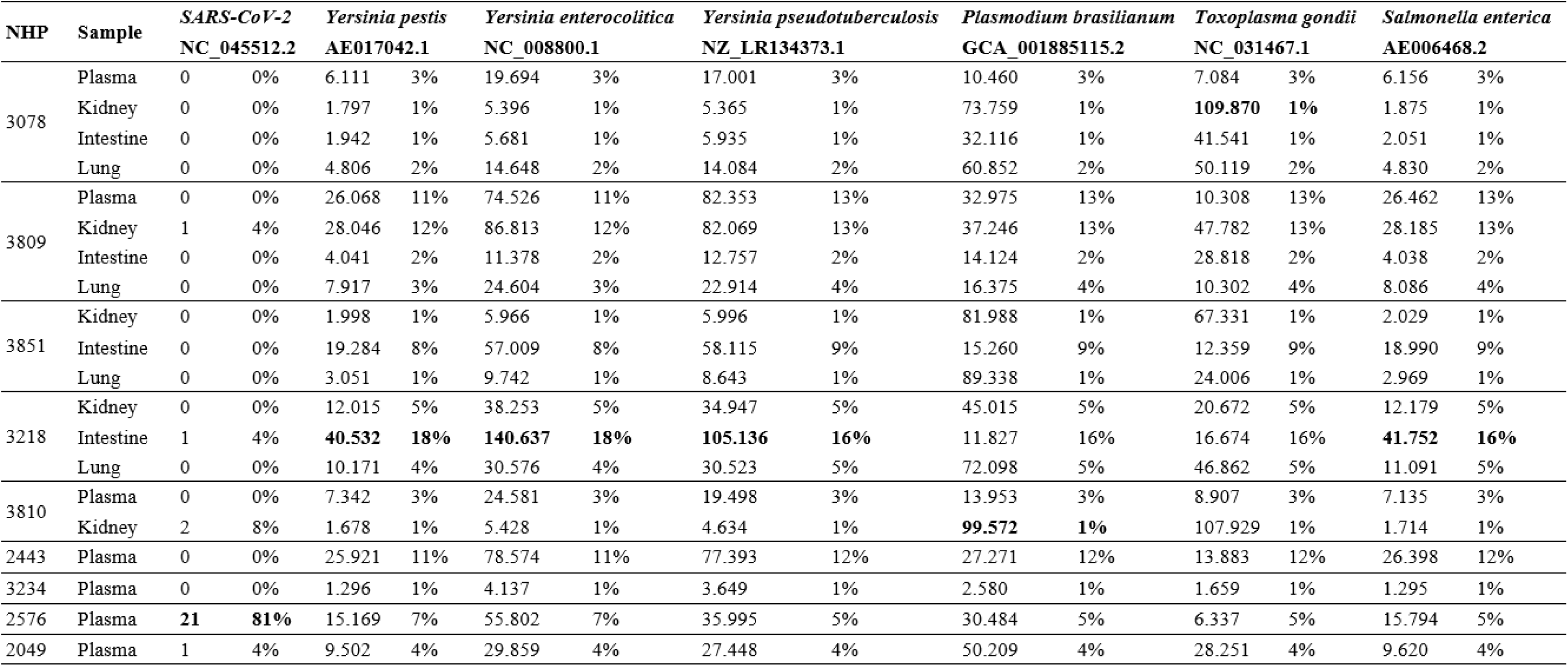
Main findings from the reference assembly using metagenomic data.

| NHP | Sample | *SARS-CoV-2* |  | *Yersinia pestis* |  | *Yersinia enterocolitica* |  | *Yersinia pseudotuberculosis* |  | *Plasmodium brasilianum* |  | *Toxoplasma gondii* |  | *Salmonella enterica* |  |
| --- | --- | --- | --- | --- | --- | --- | --- | --- | --- | --- | --- | --- | --- | --- | --- |
|  |  | NC_045512.2 |  | AE017042.1 |  | NC_008800.1 |  | NZ_LR134373.1 |  | GCA_001885115.2 |  | NC_031467.1 |  | AE006468.2 |  |
| 3078 | Plasma | 0 | 0% | 6.111 | 3% | 19.694 | 3% | 17.001 | 3% | 10.460 | 3% | 7.084 | 3% | 6.156 | 3% |
|  | Kidney | 0 | 0% | 1.797 | 1% | 5.396 | 1% | 5.365 | 1% | 73.759 | 1% | **109.870** | **1%** | 1.875 | 1% |
|  | Intestine | 0 | 0% | 1.942 | 1% | 5.681 | 1% | 5.935 | 1% | 32.116 | 1% | 41.541 | 1% | 2.051 | 1% |
|  | Lung | 0 | 0% | 4.806 | 2% | 14.648 | 2% | 14.084 | 2% | 60.852 | 2% | 50.119 | 2% | 4.830 | 2% |
| 3809 | Plasma | 0 | 0% | 26.068 | 11% | 74.526 | 11% | 82.353 | 13% | 32.975 | 13% | 10.308 | 13% | 26.462 | 13% |
|  | Kidney | 1 | 4% | 28.046 | 12% | 86.813 | 12% | 82.069 | 13% | 37.246 | 13% | 47.782 | 13% | 28.185 | 13% |
|  | Intestine | 0 | 0% | 4.041 | 2% | 11.378 | 2% | 12.757 | 2% | 14.124 | 2% | 28.818 | 2% | 4.038 | 2% |
|  | Lung | 0 | 0% | 7.917 | 3% | 24.604 | 3% | 22.914 | 4% | 16.375 | 4% | 10.302 | 4% | 8.086 | 4% |
| 3851 | Kidney | 0 | 0% | 1.998 | 1% | 5.966 | 1% | 5.996 | 1% | 81.988 | 1% | 67.331 | 1% | 2.029 | 1% |
|  | Intestine | 0 | 0% | 19.284 | 8% | 57.009 | 8% | 58.115 | 9% | 15.260 | 9% | 12.359 | 9% | 18.990 | 9% |
|  | Lung | 0 | 0% | 3.051 | 1% | 9.742 | 1% | 8.643 | 1% | 89.338 | 1% | 24.006 | 1% | 2.969 | 1% |
| 3218 | Kidney | 0 | 0% | 12.015 | 5% | 38.253 | 5% | 34.947 | 5% | 45.015 | 5% | 20.672 | 5% | 12.179 | 5% |
|  | Intestine | 1 | 4% | **40.532** | **18%** | **140.637** | **18%** | **105.136** | **16%** | 11.827 | 16% | 16.674 | 16% | **41.752** | **16%** |
|  | Lung | 0 | 0% | 10.171 | 4% | 30.576 | 4% | 30.523 | 5% | 72.098 | 5% | 46.862 | 5% | 11.091 | 5% |
| 3810 | Plasma | 0 | 0% | 7.342 | 3% | 24.581 | 3% | 19.498 | 3% | 13.953 | 3% | 8.907 | 3% | 7.135 | 3% |
|  | Kidney | 2 | 8% | 1.678 | 1% | 5.428 | 1% | 4.634 | 1% | **99.572** | **1%** | 107.929 | 1% | 1.714 | 1% |
| 2443 | Plasma | 0 | 0% | 25.921 | 11% | 78.574 | 11% | 77.393 | 12% | 27.271 | 12% | 13.883 | 12% | 26.398 | 12% |
| 3234 | Plasma | 0 | 0% | 1.296 | 1% | 4.137 | 1% | 3.649 | 1% | 2.580 | 1% | 1.659 | 1% | 1.295 | 1% |
| 2576 | Plasma | **21** | **81%** | 15.169 | 7% | 55.802 | 7% | 35.995 | 5% | 30.484 | 5% | 6.337 | 5% | 15.794 | 5% |
| 2049 | Plasma | 1 | 4% | 9.502 | 4% | 29.859 | 4% | 27.448 | 4% | 50.209 | 4% | 28.251 | 4% | 9.620 | 4% |

## Notes

### Competing Interest Statement

The authors have declared no competing interest.

## References

1. Voloch CM, da Silva Francisco R Jr, de Almeida LGP, Cardoso CC, Brustolini OJ, Gerber AL, et al. Genomic characterization of a novel SARS-CoV-2 lineage from Rio de Janeiro, Brazil. J Virol. 2021;95. doi:10.1128/JVI.00119-21

2. da Saúde S de V em S-M. PORTARIA N.° 5, DE 21 DE FEVEREIRO DE 2006. 21 Feb 2006 [cited 30 Aug 2022]. Available: https://bvsms.saude.gov.br/bvs/saudelegis/svs/2006/prt0005_21_02_2006_comp.html

3. da Saúde M. PORTARIA N° 782, DE 15 DE MARÇO DE 2017. 15 Mar 2017 [cited 16 Mar 2022]. Available: https://bvsms.saude.gov.br/bvs/saudelegis/gm/2017/prt0782_16_03_2017.html

4. da Saúde S de V em S-M. Epizootia. In: SISTEMA DE INFORMAÇÃO DE AGRAVOS DE NOTIFICAÇÃO (SINAN) [Internet]. 8 Mar 2016 [cited 17 Mar 2022]. Available: http://portalsinan.saude.gov.br/epizootia

5. de Comunicação/GAB/SVS N. Guia de vigilância de epizootias em primatas não humanos e entomologia aplicada à vigilância da febre amarela. Secretaria de Vigilância em Saúde do Ministério da Saúde; 2014. Available: https://bvsms.saude.gov.br/bvs/publicacoes/guia_vigilancia_epizootias_primatas_entomologia.pdf

6. Wada MY, Rocha SM, Maia-Elkhoury ANS. Situação da raiva no Brasil, 2000 a 2009. Epidemiologia e serviços de saúde. 2011;20: 509–518. Available: http://scielo.iec.gov.br/pdf/ess/v20n4/v20n4a10.pdf

7. Saha O, Rakhi NN, Sultana A, Rahman MM. SARS-CoV-2 and COVID-19: a threat to Global Health. Discoveries Reports. 2020;3. doi:10.15190/drep.2020.7

8. Ksiazek TG, Erdman D, Goldsmith CS, Zaki SR, Peret T, Emery S, et al. A novel coronavirus associated with severe acute respiratory syndrome. N Engl J Med. 2003;348: 1953–1966. doi:10.1056/NEJMoa030781

9. Burki T. COVID-19 in Latin America. Lancet Infect Dis. 2020;20: 547–548. doi:10.1016/S1473-3099(20)30303-0

10. Zaki AM, van Boheemen S, Bestebroer TM, Osterhaus ADME, Fouchier RAM. Isolation of a novel coronavirus from a man with pneumonia in Saudi Arabia. N Engl J Med. 2012;367: 1814– 1820. doi:10.1056/NEJMoa1211721

11. Novel Swine-Origin Influenza A (H1N1) Virus Investigation Team, Dawood FS, Jain S, Finelli L, Shaw MW, Lindstrom S, et al. Emergence of a novel swine-origin influenza A (H1N1) virus in humans. N Engl J Med. 2009;360: 2605–2615. doi:10.1056/NEJMoa0903810

12. Bell BP, Damon IK, Jernigan DB, Kenyon TA, Nichol ST, O’Connor JP, et al. Overview, Control Strategies, and Lessons Learned in the CDC Response to the 2014–2016 Ebola Epidemic. MMWR Supplements. 2016. pp. 4–11. doi:10.15585/mmwr.su6503a2

13. Slenczka WG. The Marburg virus outbreak of 1967 and subsequent episodes. Current Topics in Microbiology and Immunology. Berlin, Heidelberg: Springer Berlin Heidelberg; 1999. pp. 49–75. doi:10.1007/978-3-642-59949-1_4

14. Chang C, Ortiz K, Ansari A, Eric Gershwin M. The Zika outbreak of the 21st century. Journal of Autoimmunity. 2016. pp. 1–13. doi:10.1016/j.jaut.2016.02.006

15. Kading RC, Brault AC, Beckham JD. Global Perspectives on Arbovirus Outbreaks: A 2020 Snapshot. Trop Med Infect Dis. 2020;5. doi:10.3390/tropicalmed5030142

16. Beigel JH, Farrar J, Han AM, Hayden FG, Hyer R, de Jong MD, et al. Avian influenza A (H5N1) infection in humans. N Engl J Med. 2005;353: 1374–1385. doi:10.1056/NEJMra052211

17. European Food Safety Authority, European Centre for Disease Prevention and Control, European Union Reference Laboratory for Avian Influenza, Adlhoch C, Fusaro A, Gonzales JL, Kuiken T, Mirinaviciute G, et al. Avian influenza overview March – April 2023. EFSA J. 2023;21: e08039. doi:10.2903/j.efsa.2023.8039

18. Naranjo J, Cosivi O. Elimination of foot-and-mouth disease in South America: lessons and challenges. Philos Trans R Soc Lond B Biol Sci. 2013;368: 20120381. doi:10.1098/rstb.2012.0381

19. Shang Y, Li H, Zhang R. Effects of Pandemic Outbreak on Economies: Evidence From Business History Context. Front Public Health. 2021;9: 632043. doi:10.3389/fpubh.2021.632043

20. Alvim RGF, Lima TM, Rodrigues DAS, Marsili FF, Bozza VBT, Higa LM, et al. From a recombinant key antigen to an accurate, affordable serological test: Lessons learnt from COVID-19 for future pandemics. Biochem Eng J. 2022;186: 108537. doi:10.1016/j.bej.2022.108537

21. Hu H, Jung K, Wang Q, Saif LJ, Vlasova AN. Development of a one-step RT-PCR assay for detection of pancoronaviruses (α-, β-, γ-, and δ-coronaviruses) using newly designed degenerate primers for porcine and avian ‘fecal samples. J Virol Methods. 2018;256: 116–122. doi:10.1016/j.jviromet.2018.02.021

22. Pereira AH, Vasconcelos AL, Silva VL, Nogueira BS, Silva AC, Pacheco RC, et al. Natural SARS-CoV-2 Infection in a Free-Ranging Black-Tailed Marmoset (Mico melanurus) from an Urban Area in Mid-West Brazil. J Comp Pathol. 2022;194: 22–27. doi:10.1016/j.jcpa.2022.03.005

23. Riera LM, Feuillade MR, Saavedra MC, Ambrosio AM. Evaluation of an enzyme immunosorbent assay for the diagnosis of Argentine haemorrhagic fever. Acta Virol. 1997;41: 305–310. Available: https://www.ncbi.nlm.nih.gov/pubmed/9607087

24. Padula PJ, Rossi CM, Valle MOD, Martínez PV, Colavecchia SB, Edelstein A, et al. Development and evaluation of a solid-phase enzyme immunoassay based on Andes hantavirus recombinant nucleoprotein. J Med Microbiol. 2000;49: 149–155. doi:10.1099/0022-1317-49-2-149

25. Klempa B, Fichet-Calvet E, Lecompte E, Auste B, Aniskin V, Meisel H, et al. Hantavirus in African wood mouse, Guinea. Emerg Infect Dis. 2006;12: 838–840. doi:10.3201/eid1205.051487

26. Guterres A, de Oliveira RC, Fernandes J, Schrago CG, de Lemos ERS. Detection of different South American hantaviruses. Virus Res. 2015;210: 106–113. doi:10.1016/j.virusres.2015.07.022

27. García JB, Morzunov SP, Levis S, Rowe J, Calderón G, Enría D, et al. Genetic diversity of the Junin virus in Argentina: geographic and temporal patterns. Virology. 2000;272: 127–136. doi:10.1006/viro.2000.0345

28. Emonet S, Retornaz K, Gonzalez J-P, de Lamballerie X, Charrel RN. Mouse-to-human transmission of variant lymphocytic choriomeningitis virus. Emerg Infect Dis. 2007;13: 472–475. doi:10.3201/eid1303.061141

29. Domingo C, Patel P, Yillah J, Weidmann M, Méndez JA, Nakouné ER, et al. Advanced yellow fever virus genome detection in point-of-care facilities and reference laboratories. J Clin Microbiol. 2012;50: 4054–4060. doi:10.1128/JCM.01799-12

30. Lanciotti RS, Kosoy OL, Laven JJ, Velez JO, Lambert AJ, Johnson AJ, et al. Genetic and serologic properties of Zika virus associated with an epidemic, Yap State, Micronesia, 2007. Emerg Infect Dis. 2008;14: 1232–1239. doi:10.3201/eid1408.080287

31. Ribeiro MO, Godoy DT, Fontana-Maurell M, Costa EM, Andrade EF, Rocha DR, et al. Analytical and clinical performance of molecular assay used by the Brazilian public laboratory network to detect and discriminate Zika, Dengue and Chikungunya viruses in blood. Braz J Infect Dis. 2021;25: 101542. doi:10.1016/j.bjid.2021.101542

32. Lanciotti RS, Kosoy OL, Laven JJ, Panella AJ, Velez JO, Lambert AJ, et al. Chikungunya virus in US travelers returning from India, 2006. Emerg Infect Dis. 2007;13: 764–767. doi:10.3201/eid1305.070015

33. de Paula Silveira-Lacerda E, Laschuk Herlinger A, Tanuri A, Rezza G, Anunciação CE, Ribeiro JP, et al. Molecular epidemiological investigation of Mayaro virus in febrile patients from Goiania City, 2017-2018. Infect Genet Evol. 2021;95: 104981. doi:10.1016/j.meegid.2021.104981

34. Lanciotti RS, Kerst AJ, Nasci RS, Godsey MS, Mitchell CJ, Savage HM, et al. Rapid detection of west nile virus from human clinical specimens, field-collected mosquitoes, and avian samples by a TaqMan reverse transcriptase-PCR assay. J Clin Microbiol. 2000;38: 4066–4071. doi:10.1128/JCM.38.11.4066-4071.2000

35. Naveca FG, Nascimento VA do, Souza VC de, Nunes BTD, Rodrigues DSG, Vasconcelos PF da C. Multiplexed reverse transcription real-time polymerase chain reaction for simultaneous detection of Mayaro, Oropouche, and Oropouche-like viruses. Mem Inst Oswaldo Cruz. 2017;112: 510–513. doi:10.1590/0074-02760160062

36. Claro IM, Ramundo MS, Coletti TM, da Silva CAM, Valenca IN, Candido DS, et al. Rapid viral metagenomics using SMART-9N amplification and nanopore sequencing. Wellcome Open Res. 2021;6: 241. doi:10.12688/wellcomeopenres.17170.1

37. Wood DE, Lu J, Langmead B. Improved metagenomic analysis with Kraken 2. Genome Biol. 2019;20: 257. doi:10.1186/s13059-019-1891-0

38. Ondov BD, Bergman NH, Phillippy AM. Interactive metagenomic visualization in a Web browser. BMC Bioinformatics. 2011;12: 385. doi:10.1186/1471-2105-12-385

39. Li H. Minimap2: pairwise alignment for nucleotide sequences. Bioinformatics. 2018;34: 3094–3100. doi:10.1093/bioinformatics/bty191

40. Milne I, Bayer M, Stephen G, Cardle L, Marshall D. Tablet: Visualizing Next-Generation Sequence Assemblies and Mappings. In: Edwards D, editor. Plant Bioinformatics: Methods and Protocols. New York, NY: Springer New York; 2016. pp. 253–268. doi:10.1007/978-1-4939-3167-5_14

41. Giles J, Peterson AT, Almeida A. Ecology and geography of plague transmission areas in northeastern Brazil. PLoS Negl Trop Dis. 2011;5: e925. doi:10.1371/journal.pntd.0000925

42. Bezerra MF, Xavier CC, Almeida AMP de, Reis CR de S. Evaluation of a multi-species Protein A-ELISA assay for plague serologic diagnosis in humans and other mammal hosts. PLoS Negl Trop Dis. 2022;16: e0009805. doi:10.1371/journal.pntd.0009805

43. Chu MC. Laboratory Manual of Plague Diagnostic Tests. Center for Disease Control and Prevention (CDC); 2000. Available: https://play.google.com/store/books/details?id=_9wMtAEACAAJ

44. Leal NC, Almeida AM. Diagnosis of plague and identification of virulence markers in Yersinia pestis by multiplex-PCR. Rev Inst Med Trop Sao Paulo. 1999;41: 339–342. doi:10.1590/s0036-46651999000600002

45. Konradt G, Bassuino DM, Prates KS, Bianchi MV, Snel GGM, Sonne L, et al. Suppurative infectious diseases of the central nervous system in domestic ruminants. Pesqui Vet Bras. 2017;37: 820–828. doi:10.1590/S0100-736X2017000800007

46. Pereira GO, Pereira AHB, Brito M de F, Pescador CA, Ubiali DG. Toxoplasma gondii induced abortions in a goat herd in Rio de Janeiro, Brazil. Cienc Rural. 2021;51. doi:10.1590/0103-8478cr20200568

47. Villar-Echarte G, Arruda IF, Barbosa A da S, Guzmán RG, Augusto AM, Troccoli F, et al. Toxoplasma gondii among captive wild mammals in zoos in Brazil and Cuba: seroprevalence and associated risk factors. Rev Bras Parasitol Vet. 2021;30: e001921. doi:10.1590/S1984-29612021053

48. Homan WL, Vercammen M, De Braekeleer J, Verschueren H. Identification of a 200– to 300-fold repetitive 529 bp DNA fragment in Toxoplasma gondii, and its use for diagnostic and quantitative PCR. Int J Parasitol. 2000;30: 69–75. doi:10.1016/s0020-7519(99)00170-8

49. Hall T, Biosciences I, Carlsbad C, Others. BioEdit: an important software for molecular biology. GERF Bull Biosci. 2011;2: 60–61. Available: https://www.academia.edu/download/32793732/Bioedit_software_review.pdf

50. Moreira SB, Pereira AHB, Pissinatti T de A, Arruda IF, de Azevedo RRM, Schiffler FB, et al. Subacute multisystemic toxoplasmosis in a captive black-and-gold howler monkey (Alouatta caraya) indicates therapy challenging. J Med Primatol. 2022;51: 392–395. doi:10.1111/jmp.12600

51. Dobson AP, Robin Carper E. Infectious Diseases and Human Population History. BioScience. 1996. pp. 115–126. doi:10.2307/1312814

52. Levinson J, Bogich T, Olival K, Epstein J, Johnson C, Karesh W, et al. Targeting Surveillance for Zoonotic Virus Discovery. Emerging Infectious Disease journal. 2013;19: 743. doi:10.3201/eid1905.121042

53. Kowalewski MM, Salzer JS, Deutsch JC, Raño M, Kuhlenschmidt MS, Gillespie TR. Black and gold howler monkeys (Alouatta caraya) as sentinels of ecosystem health: patterns of zoonotic protozoa infection relative to degree of human-primate contact. Am J Primatol. 2011;73: 75–83. doi:10.1002/ajp.20803

54. Batista PM. Arbovírus em primatas não humanos capturados em Mato Grosso do Sul. PhD, Fundação Universidade Federal de Mato Grosso do Sul. 2015. Available: https://repositorio.ufms.br/handle/123456789/2918

55. Rocha TC da, Batista PM, Andreotti R, Bona ACD, Silva MAN da, Lange R, et al. Evaluation of arboviruses of public health interest in free-living non-human primates (*Alouatta* spp., *Callithrix* spp., Sapajus spp.) in Brazil. Rev Soc Bras Med Trop. 2015;48: 143–148. doi:10.1590/0037-8682-0024-2015

56. Santos SV, Pena HFJ, Talebi MG, Teixeira RHF, Kanamura CT, Diaz-Delgado J, et al. Fatal toxoplasmosis in a southern muriqui (Brachyteles arachnoides) from São Paulo state, Brazil: Pathological, immunohistochemical, and molecular characterization. Journal of Medical Primatology. 2017;47: 124–127. doi:10.1111/jmp.12326

57. Kompalic-Cristo A, Britto C, Fernandes O. Diagnóstico molecular da toxoplasmose: revisão. J Bras Patol Med Lab. 2005;41: 229–235. doi:10.1590/S1676-24442005000400003

58. Frenkel JK, Dubey JP, Beattie CP. Toxoplasmosis of Animals and Man. J Parasitol. 1989;75: 816. doi:10.2307/3283074

59. Dubey JP. Toxoplasmosis of Animals and Humans. 2nd Edition. CRC Press; 2009. doi:10.1201/9781420092370

60. Dubey JP, Murata FHA, Cerqueira-Cézar CK, Kwok OCH, Yang Y, Su C. Recent epidemiologic, clinical, and genetic diversity of Toxoplasma gondii infections in non-human primates. Res Vet Sci. 2021;136: 631–641. doi:10.1016/j.rvsc.2021.04.017

61. Henrik Dietz H, Henriksen P, Bille-Hansen V, Aage Henriksen S. Toxoplasmosis in a colony of New World monkeys. Vet Parasitol. 1997;68: 299–304. doi:10.1016/S0304-4017(96)01088-6

62. Bouer A, Werther K, Catão-Dias JL, Nunes ALV. Outbreak of Toxoplasmosis in Lagothrix lagotricha. Folia Primatol. 1999;70: 282–285. doi:10.1159/000021709

63. Epiphanio S, Guimarães MAB, Fedullo DL, Correa SHR, Catão-Dias JL. Toxoplasmosis in Golden-headed Lion Tamarins (Leontopithecus chrysomelas) and Emperor Marmosets (Saguinus imperator) in captivity. zamd. 2000;31: 231–235. doi:10.1638/1042-7260(2000)031[0231:TIGHLT]2.0.CO;2

64. Epiphanio S, Sá LR, Teixeira RH, Catão-Dias JL. Toxoplasmosis in a wild-caught black lion tamarin (Leontopithecus chrysopygus). Vet Rec. 2001;149: 627–628. doi:10.1136/vr.149.20.627

65. Epiphanio S, Sinhorini IL, Catão-Dias JL. Pathology of toxoplasmosis in captive new world primates. J Comp Pathol. 2003;129: 196–204. doi:10.1016/s0021-9975(03)00035-5

66. Casagrande RA, Silva TCE da, Pescador CA, Borelli V, Souza JC Jr, Souza ER, et al. Toxoplasmose em primatas neotropicais: estudo retrospectivo de sete casos. Pesqui Vet Bras. 2013;33: 94–98. doi:10.1590/S0100-736X2013000100017

67. Reis Amendoeira MR, Arruda IF, Moreira SB, Ubiali DG, da Silva Barbosa A, Jesus Pena HF, et al. Isolation and genetic characterization of Toxoplasma gondii from a captive black-and-gold howler monkey (Alouatta caraya Humboldt, 1812) in Brazil. Int J Parasitol Parasites Wildl. 2022;19: 187–190. doi:10.1016/j.ijppaw.2022.09.005

68. Pena HFJ, Vitaliano SN, Beltrame MAV, Pereira FEL, Gennari SM, Soares RM. PCR-RFLP genotyping of Toxoplasma gondii from chickens from Espírito Santo state, Southeast region, Brazil: new genotypes and a new SAG3 marker allele. Vet Parasitol. 2013;192: 111–117. doi:10.1016/j.vetpar.2012.10.004

69. Ferreira TCR, Buery JC, Moreira NIB, Santos CB, Costa JGL, Pinto LV, et al. Toxoplasma gondii: isolation, biological and molecular characterisation of samples from free-range Gallus gallus domesticus from countryside Southeast Brazil. Rev Bras Parasitol Vet. 2018;27: 384–389. doi:10.1590/s1984-296120180028

70. Silva LA, Andrade RO, Carneiro ACAV, Vitor RWA. Overlapping Toxoplasma gondii genotypes circulating in domestic animals and humans in Southeastern Brazil. PLoS One. 2014;9: e90237. doi:10.1371/journal.pone.0090237

71. Carneiro ACAV, Andrade GM, Costa JGL, Pinheiro BV, Vasconcelos-Santos DV, Ferreira AM, et al. Genetic characterization of Toxoplasma gondii revealed highly diverse genotypes for isolates from newborns with congenital toxoplasmosis in southeastern Brazil. J Clin Microbiol. 2013;51: 901–907. doi:10.1128/JCM.02502-12

72. Paula NF de, Dutra KS, Oliveira AR de, Santos DOD, Rocha CEV, Vitor RW de A, et al. Host range and susceptibility to Toxoplasma gondii infection in captive neotropical and Old-world primates. J Med Primatol. 2020;49: 202–210. doi:10.1111/jmp.12470

73. de Vigilância das Arboviroses (CGARB) C-G. Febre Amarela em humanos e primatas não-humanos – 1994 a 2021 – OPENDATASUS. 2022. Available: https://opendatasus.saude.gov.br/dataset/febre-amarela-em-humanos-e-primatas-nao-humanos

74. Calderon C, Moreira A, Marquez ES. Herpersvirusis in non-human primates. Scientific Electronic. 2016. Available: https://sea.ufr.edu.br/SEA/article/view/283/

75. IUCN. The IUCN Red List of Threatened Species. Version 2022-2. In: IUCN Red List of Threatened Species [Internet]. [cited 22 Mar 2023]. Available: https://www.iucnredlist.org

